# A variational deep-learning approach to modeling memory T cell dynamics

**DOI:** 10.1101/2024.07.08.602409

**Authors:** Christiaan H. van Dorp, Joshua I. Gray, Daniel H. Paik, Donna L. Farber, Andrew J. Yates

**Author notes:** Authors contributed equally (Christiaan H. van Dorp, Joshua I. Gray).

## Abstract

Mechanistic models of dynamic, interacting cell populations have yielded many insights into the growth and resolution of immune responses. Historically these models have described the behavior of pre-defined cell types based on small numbers of phenotypic markers. The ubiquity of deep phenotyping therefore presents a new challenge; how do we confront tractable and interpretable mathematical models with high-dimensional data? To tackle this problem, we studied the development and persistence of lung-resident memory CD4 and CD8 T cells (T_RM_) in mice infected with influenza virus. We developed an approach in which dynamical model parameters and the population structure are inferred simultaneously. This method uses deep learning and stochastic variational inference and is trained on the single-cell flow-cytometry data directly, rather than on the kinetics of pre-identified clusters. We show that during the resolution phase of the immune response, memory CD4 and CD8 T cells within the lung are phenotypically diverse, with subsets exhibiting highly distinct and time-dependent dynamics. T_RM_ heterogeneity is maintained long-term by ongoing differentiation of relatively persistent Bcl-2hi CD4 and CD8 T_RM_ subsets which resolve into distinct functional populations. Our approach yields new insights into the dynamics of tissue-localized immune memory, and is a novel basis for interpreting time series of high-dimensional data, broadly applicable to diverse biological systems.

## Introduction

The widespread implementation of single-cell high-throughput assays has driven the development of a plethora of computational tools for generating lower-dimensional (‘latent’) representations of datasets that occupy high-dimensional ‘feature’ spaces, usually involving the identification of discrete clusters of cellular states or phenotypes [1, 2]. Typically, inferences are derived from single snapshots of systems at steady state. However, following the response of a system to perturbations has the potential to reveal more about the components and interactions that underpin it. Mathematical models are used widely for formulating mechanistic descriptions of cellular dynamics, and drawing inferences from time-series of observations. Such models usually describe the directly-observed trajectories of small numbers of discrete, pre-defined cell types or states. An outstanding challenge, then, is to connect dynamical modeling tools to exploit the information contained with the burgeoning quantities of high dimensional data that are now available.

An acute immune response is an archetype of a dynamic, perturbed set of biological processes. In particular, T cells that respond to infectious challenge undergo clonal expansion and diversification, generating heterogeneous populations of memory-phenotype cells that are capable of responding rapidly and potently to re-exposure to the same infectious agent. These memory T cells can be broadly characterized as circulating or tissue-resident [3]. To date, the majority of modeling studies have focused on the dynamics of circulating memory subsets in both humans and mouse models, characterizing their ontogeny, diversity, kinetics, and persistence [4–8]. In contrast, the dynamics of tissue-resident memory T cells (T_RM_), which patrol organs and barrier sites and provide indispensable front-line protection against a wide variety of pathogens [9], are relatively understudied [10]. Understanding how T_RM_ are established and maintained, and what determines the duration of protection that they offer, has the potential to revolutionize vaccine design [11], our understanding of autoimmune responses, and the control of tumor growth.

As with cell types in many areas of biology, T_RM_ are usually defined by combinations of surface markers and are traditionally identified with flow cytometry using hierarchical binary gating strategies [11]. CD8 T_RM_ in the lung are commonly associated with CD103, which binds to E-cadherin present on epithelial cells, supporting retention in barrier tissues, and CD69, which limits tissue egress by antagonizing S1P1-mediated extravasation [9, 11]. CD4 T_RM_ in the lung are typically characterized by expression of CD69 and the chemokine receptor CXCR6 [12, 13]. However, the expression patterns of many of these markers do not yield objectively defined cutoff values to characterize positive and negative populations unambiguously, a problem compounded as the number of these markers grows [14]. The ontogeny and persistence of T_RM_ therefore provide an ideal setting to explore the application of dynamical modeling to high-dimensional phenotyping data.

An obvious approach is to perform initial unsupervised clustering step, which combines dimension reduction and phenotypic classification. One can then use sets of ordinary differential equations (ODEs) or other dynamical systems to describe the time evolution of the sizes of these clusters [15], some of which may represent phenotypes that are not canonically defined. This process typically yields estimates of rates of cell loss, self-renewal, differentiation, and cell killing. In principle this sequential approach is highly tractable, because many off-the-shelf clustering methods are available, and standard tools can be applied to perform inference and model selection. Potential problems remain, however. First, clusters are usually not clearly separable and so in general the assignment of cells to clusters should be probabilistic. Second, as the phenotypic structure of a population that is out of equilibrium evolves, the uncertainty in cluster assignment may also be time dependent. These issues suggest that under some conditions it may be necessary to jointly model the distribution of the flow cytometry data itself, as well as the underlying cellular dynamics.

To explore these issues, we used high dimensional flow cytometry to characterize the dynamics and phenotypic structure of T cell subsets within the lungs of mice that were infected with influenza A virus (IAV) and followed up to 57 days post-infection. To model the time evolution of these populations, we developed and compared two methodologies. The first, ‘sequential’ approach employs standard clustering methods applied to the processed flow cytometry data pooled across time points, and models the time evolution of the cluster sizes with ODE models. The second, ‘integrated’ method employs deep learning and stochastic variational inference [16] to simultaneously model the structure and dynamics of observed marker expression, via a lower dimensional representation of the data. With these approaches we find that both CD4 and CD8 T cells within the lung are phenotypically and dynamically heterogeneous. We also identified persistent populations within both subsets that may maintain this diversity over weeks to months through continuous differentiation. We discuss the benefits and limitations of the two approaches to modeling high dimensional time series, and conclude by highlighting areas for further development.

## Results

### The size and phenotypic composition of lung-localized T cell populations change dynamically following influenza infection in mice

The mouse model of influenza infection recapitulates the dynamics of virus morbidity and the immune response seen in human infection, and results in the generation of lung CD4 and CD8 tissue resident memory T cells [17, 18] (T_RM_). We followed the T cell responses to influenza A virus (IAV) infection in mice, at timepoints between 6 and 57 days post-infection (DPI; Fig. 1A; see Methods). We restricted our analyses to antigen-experienced (CD44+ CD11a+) CD8+ and CD4+ T cells (which hereon we refer to as CD8 and CD4 T cells, for brevity) within the lung tissue, as defined by protection from *in vivo* antibody labeling as described [17, 19] (Fig. S1). We previously determined that T cells in this lung resident niche represent the majority of influenza-specific memory T cells [19]. Both populations peaked in numbers between 8-10 DPI, shortly after the point of maximum weight loss as a measure of infection morbidity (Fig. S2A), and then declined with population half-lives of roughly 3 and 5 days respectively (Fig. 1B and C). Both CD8 and CD4 T cells approached stable numbers by 37 DPI, after contracting approximately 100-fold and 10-fold, respectively.

**Figure 1:**
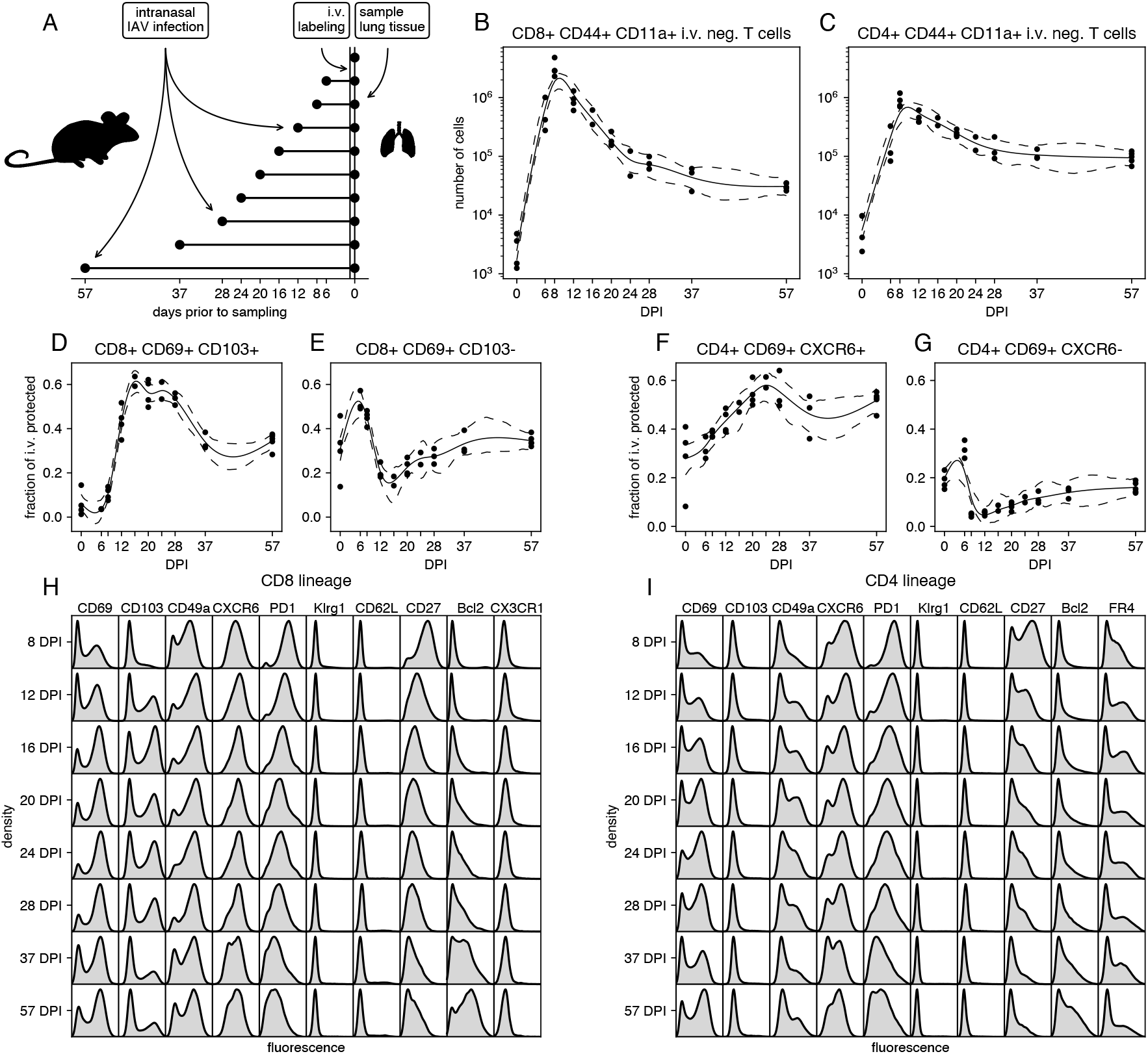
Studying the timecourse of the lung-localized CD8 and CD4 T cell responses to IAV infection. **A**. Experimental design. Each cohort contained 4 or 5 mice and 42 mice were used in total. Eight mice were excluded due to lack of weight loss or i.v. label failure (Fig S2). **B-C**. Total numbers of antigen-experienced, protected CD8+ and conventional CD4+ T cells by day post infection (DPI), calculated by multiplying the estimated total number of lung cells with the fraction of protected CD44+ CD11a+ CD8+ and conventional CD4+ T cells. **D-E**. Frequencies of cells expressing combinations of the tissue-resident memory markers CD69 and CD103 within protected CD44+ CD11a+ CD8+ T cells. **F-G**. Frequencies of cells expressing combinations of the tissue-resident memory markers CD69 and CXCR6 within protected CD44+ CD11a+ CD4+ T cells. **H-I**. Marginal distributions of expression of a selection of markers, aggregated over mice and stratified by day post infection, and lineage (CD8 or CD4).

These smooth trajectories were highly characteristic of adaptive immune responses and of IAV responses in particular [20, 21]. However, within them, we identified considerable heterogeneity with respect to surface markers of tissue residency [11, 18, 20]. The (relative) abundances of T cell subsets defined by combinations of the canonical markers CD69, CD103 and CXCR6 clearly shifted during the contraction phase of the response (Figs. 1D-G). For example, the relative abundance of CD8 T cells expressing both CD69 and CD103 increased with time, then decreased, while CD8 T cells expressing CD69 but not CD103 were rapidly lost in the second week post infection, and then stabilized (Fig. 1D-E). Around half of CD4 T cells expressed both CD69 and CXCR6 beyond 4 weeks post infection. CD4 T cells with low CXCR6 and high CD69 expression mostly disappeared around the peak of the immune response, and then very slowly grew in proportion (Fig. 1F-G). This (temporal) heterogeneity was manifested more clearly when phenotyping was expanded to include markers of cell survival (Bcl-2), access to lymphoid organs (CD62L, typically used to define circulating ‘central’ memory cells, T_CM_), differentiation (KLRG1 and PD-1), a costimulatory receptor associated with cell survival (CD27), the chemokine receptor CX3CR1, the follicular helper T cell marker FR4, and the integrin CD49a. The marginal distributions of the expression of many of these markers showed clear variation with time (Fig. 1H and I). Thus, both CD4 and CD8 T cells within the lung exhibit complex and shifting phenotypic structures during the contraction and memory phases of IAV infection.

### Sequential Approach

#### Standard unbiased clustering methods clearly delineate heterogeneity within CD8 T cells

To understand and quantify how these structures emerge and evolve, we began with a sequential approach to modeling the population dynamics of T cells within the lung from the peak of infection onward. For clarity, we first focus largely on the CD8 T cell response, and then present a more condensed description of the CD4 T cell response that highlights the key results.

We aggregated the flow cytometric data from all mice and all time points, and then assigned CD8 T cells to populations using the Leiden clustering method [22] and manual annotation. For each mouse, we tabulated the proportions of cells assigned to each population. Together with the measured total numbers of antigen-experienced protected CD8 T cells in the lung, this process yielded time series of the cluster sizes as a function of time post infection. We used Bayesian dynamical models and the leave-one-out information criterion, implemented in Stan [23, 24], to infer biological parameters from these time series and assess the relative support for different models. The workflow of the sequential approach is summarized in Fig. 2A.

**Figure 2:**
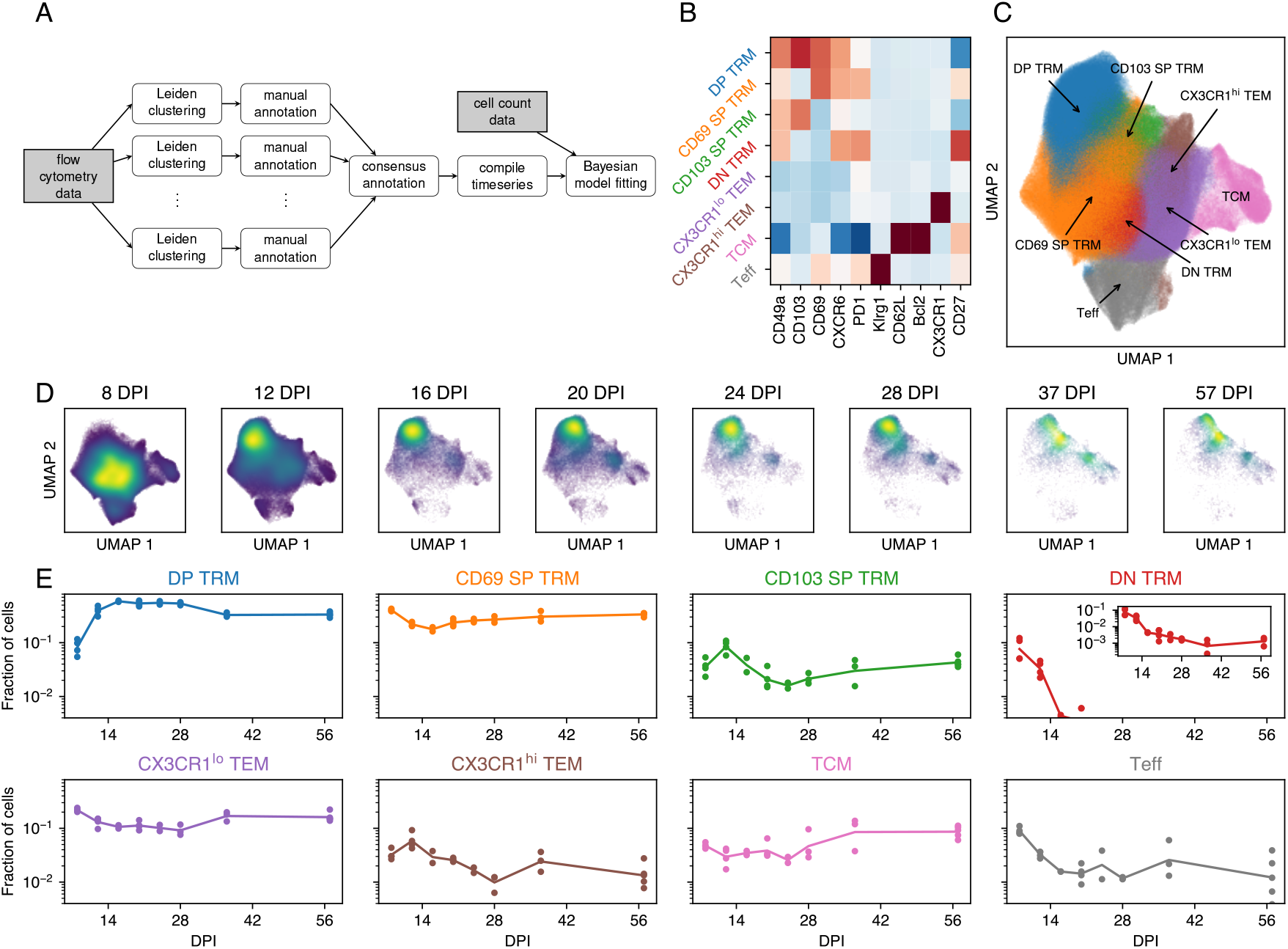
Pre-processing flow cytometry data for the sequential approach. Results are based on data from *n* = 27 mice. **A**.Flow-chart of the sequential approach. **B**. Marker expression heatmap for selected markers and consensus T-cell populations. **C**. Uniform Manifold Approximation and Projection [25] embedding (UMAP) of the marker expression data, colored by annotation. **D**. UMAPs of marker expression data, split by day post infection (DPI). The color scale reflects cell density in UMAP space. **E**. Time series of the fraction of cells in each cluster. The lines show a linear interpolation on the log scale. The panels have the same *y*-scale for easy comparison and the inset axis shows the lower values of the DN T_RM_ population.

The Leiden algorithm is stochastic, and we found that both the number and marker expression profiles of the clusters varied from run to run. To address this uncertainty we performed a consensus clustering step. We ran the Leiden method 20 times, and manually annotated the clusters as cell types. Each cell’s type was determined by its consensus assignment. We return to the issue of uncertainty in cluster assignment below.

The Leiden clustering robustly identified eight distinct CD8 T cell populations (Fig. 2B and Fig. 2C). Central memory T cells (T_CM_) were CD62Lhi Bcl-2hi, and an effector-like population (T_Eff_) was clearly defined by high levels of expression of KLRG1. We identified four T_RM_-like populations based on expression of CD69 and CD103, together with CD49a, CXCR6, and PD-1. We denoted them DP T_RM_ (CD69hi CD103hi), CD69 SP T_RM_ (CD69hi CD103lo), CD103 SP T_RM_ (CD69lo CD103hi), and DN T_RM_ (CD69lo CD103lo), but note that these populations also vary in their expression of other markers. We also identified a population which we define as effector memory (T_EM_) cells, which lacked expression of markers conventionally associated with T_CM_ and T_RM_. A subset of these T_EM_ cells expressed CX3CR1 at high levels, which has been postulated to be a key marker for defining memory CD8 T cell subsets [26] and may indicate a reduced capacity for residency [27].

#### IAV nucleoprotein-specific T cells in lung cluster similarly to their polyclonal parent populations

To further validate our clustering step, we used a fluorescently labeled MHC class I tetramer to identify CD8 T cells specific for the NP_311-325_ epitope of IAV, and examined their distribution relative to the cluster locations identified for the response in total (Fig. S3). The tetramer-positive (Tet+) populations represented approximately 10% of CD8 T cells in bulk, and exhibited similar kinetics (Fig. S4), in concordance with previous findings [19]. The partitioning of the UMAP space into 8 sub-populations appeared to respect the density of Tet+ cells (Fig. S3), although they were somewhat enriched within the CD69 SP T_RM_, DN T_RM_ and both T_EM_ populations, suggesting some variation in phenotypic structure with epitope specificity, and/or that a proportion of CD8 T cells within the lung were not IAV-specific. Similar observations held for the NP_311-325_-specific CD4 T cells, identified with an MHC class II tetramer, which represented roughly 5% of total CD4 and were slightly enriched for an effector-like population (Fig. S3). Overall, however, the congruent kinetics of epitope-specific and bulk populations, the fact that these bulk lung niche cells are likely IAV-specific [19], and the improved resolution provided by greater cell numbers, lead us to continue with our analyses of the bulk lung niche CD4 and CD8 T cells.

#### Dynamical modeling reveals evidence for ongoing differentiation and time-dependent loss rates of lung CD8 T cell populations

Having partitioned the CD8 T cells among the eight populations, we visualized their time evolution qualitatively by UMAP (Fig. 2D) and quantitatively, showing their abundances within the population as a whole (Fig. 2E). Overall, the dominant populations were DP T_RM_, CD69 SP T_RM_ and T_EM_, with a relatively stable but lower representation (5-10%) of T_CM_. The T_Eff_ proportion quickly fell during the resolution phase to stabilize at around 1% by 16 DPI, while the DN T_RM_ fraction became almost undetectable over the same timescale. The dynamics of the DP T_RM_ and CD69 SP T_RM_ proportions resembled those derived by manual gating (Fig. 1D and E), serving as additional validation of the Leiden clustering approach.

To quantify the cellular dynamics underlying these shifts in phenotype, we fitted several models to the timecourses of cluster sizes (Fig. 3A). In the simplest model (I), we estimated a single net loss rate (the balance of loss due to cell death and/or egress from the lung, and self-renewal) for each population. In model II we allowed the net loss rates to be time-dependent, reflecting for example changes in inflammation level within the lung after the virus is cleared. In more flexible models we also allowed differentiation between populations, both without (model III) and with (model IV) time-dependent loss. To quantify the effect of migration of circulating T cells to the lung during the memory contraction phase, we performed a cell transfer experiment using CD90 congenic mice (Fig. S5 and SI Text A), which suggested that such ingress contributes minimally to the population dynamics after the infection is cleared. In all models we therefore make the simplifying assumption that ingress is negligible.

**Figure 3:**
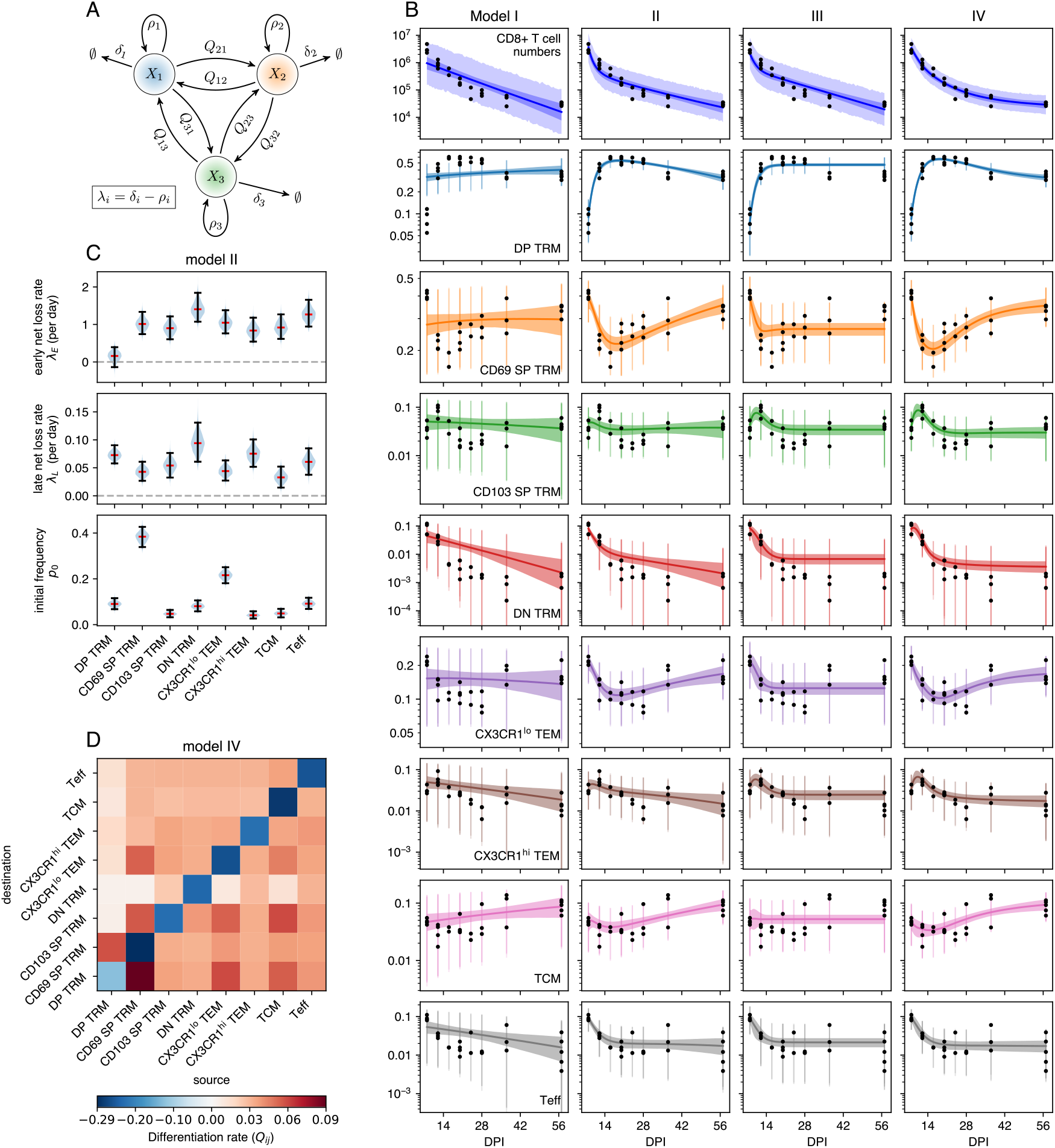
Modeling lung resident CD8 T cell dynamics using the sequential approach. Results are based on data from *n* = 27 mice. **A**. Schematic of the mathematical models, illustrated with 3 populations. *X*_*i*_ is the number of cells in population *i* and *λ*_*i*_ = *δ*_*i*_ − *ρ*_*i*_ its (possibly time dependent) net loss rate. *Q*_*ji*_ is the *per capita* rate of differentiation from *i* to *j*. **B**. Top panels: numbers of CD8+ T cells in the lung (black dots) and the model fits (blue lines). Remaining panels: observed and predicted subpopulation frequencies. Bands indicate 95% credible envelopes, bars indicate the 2.5 and 97.5 percentiles of the posterior predictive distributions. **C**. For model II, marginal posterior distributions of the parameters as violin plots (blue), with median (red) and 95% credible intervals (CrI; black). **D**. Differentiation rates in model IV. Diagonal; total egress rate by differentiation.

#### The loss of lung-resident CD8 T cells slows progressively following the resolution of IAV infection

From 8 DPI, the biphasic loss of CD8 T cells in the lung (Fig. 1C) could in principle be described by model I, in which two or more populations are lost at different but constant rates (SI Text B). A general feature of this model is that the trajectories of the subpopulation frequencies on a logarithmic scale are necessarily linear or concave, but not convex (SI Text C). The data were clearly incompatible with this behavior (Fig. 3B), and so we rejected it. In model II we allowed the rate of loss of each population to transition between initial and longer-term values at an exponential rate *u*. For simplicity, we assumed that this modulation in dynamics applied to all populations in tandem, potentially reflecting global changes in the lung environment as inflammation is resolved. This model yielded a markedly improved fit (Fig. 3B, second column), allowing for convexity in population frequencies (notably CD69 SP T_RM_, T_EM_, and T_Eff_), and better capturing the biphasic decline in total cell numbers. The estimated timescale of transition between the early and long-term dynamics, 1*/u*, was approximately 3 days, reflecting the large shifts in relative abundances of phenotypes between 8 and 14 DPI. Formal comparison of models I and II using LOO-IC [28] confirmed strong support for time-dependent loss rates (ΔLOO-IC = 130 *±* 31; see Table S1). The long-term loss rates *λ*_*L*_ yielded by model II, shown in Fig. 3C, are consistent with our prior estimates that influenza-specific CD8 T cells are lost from the lung with a population half life (ln(2)*/λ*_*L*_; reflecting the net effect of loss and any self-renewal) of 18-25 days [10].

#### Evidence for ongoing differentiation of CD8 T cell subsets within the lung

Model III extended model I to allow differentiation between CD8 phenotypes after 8 DPI. Allowing differentiation between every pair of populations risks overfitting, and so we penalized large values of *Q*_*ij*_ by regularizing them with an exponential prior distribution, similar to the lasso [29]. A key prediction of model III is that it allows subset frequencies to stabilize at late time points, as was observed for some populations (Fig. 3B, third column), because the balance of differentiation and loss can generate stable equilibria. Indeed we found substantial support for differentiation (model III) over the simplest model with constant loss rates, model I (ΔLOO-IC = 82 *±* 29). However, time-dependent loss only (model II) was still supported most strongly. We then explored model IV that combined the features of models II and III. This model was able to capture both the rapid early dynamics and long-term stability of subpopulation frequencies (Fig. 3B, fourth column), and it was very strongly supported statistically (ΔLOO-IC = 29 *±* 6.8, relative to model II). Indeed, after translating LOO-IC to Bayesian model weights [30], model IV had close to 100% support among the four candidates. However, it predicted differentiation between nearly all subpopulations (Fig. 3D), at rates with wide credible intervals (not shown). Further, the net loss rates had strong posterior correlations with the differentiation rates, indicating that the model can trade off loss with differentiation. For example, high rates of death and/or onward differentiation, balanced by influx from a precursor, are difficult to discriminate from a more isolated population that is terminally differentiated. This tension between quality of fit and interpretability highlights the limitations of relying on statistical support alone to select models.

#### Parameter and model identifiability

The potential trade-off between differentiation and loss described above led us to explore whether the parameters of the model can be identified given the available data. As a first step, we performed a structural identifiably analysis (Methods), which revealed that with perfect data (*i*.*e*. unlimited replicates and dense in time), all model parameters are globally identifiable. Our data are not perfect in this sense, and hence we also explored parameter sensitivity and practical identifiability.

The identifiability of a parameter can be assessed by changing its value by a small amount and observing the changes in the system trajectories (SI Text E). If none of the trajectories are sensitive to changes in a parameter at any time point, then estimating this parameter reliably may be difficult. We focused on the sensitivity of the trajectories *π*_*k*_(*t*) and *Y* (*t*) to perturbations of the differentiation rates *Q*_*ij*_ (Fig. S12AB and S13AB). As there are many combinations of *π*_*k*_ and *Q*_*ij*_, we focused on direct effects: How does changing influx (*Q*_*ij*_, black curves) or efflux (*Q*_*ji*_, red curves) influence the trajectory *π*_*i*_ of T-cell population *i*? As expected, the sensitivity of a trajectory *π*_*k*_ where *k ≠ i, j* to a differentiation rate *Q*_*ij*_ is generally smaller than the direct effects (Fig. S12A and S13A gray curves). Not surprisingly, increasing the influx from another population increases the relative frequency of the target population, and *vice versa*. However, these effects are most noticeable when the source population is large compared to the target (DP TRM, CD69 SP TRM, CX3CR1lo TEM columns of Fig. S12A). This effect can also depend on time. For instance, Bcl-2hi CD4 T_RM_ dominate at late time points, and the trajectories *π*_*i*_(*t*) of other populations *i* are highly sensitive to the rate of influx from Bcl-2hi T_RM_ as *t* approaches 57 DPI (Fig S13A, Bcl-2hi TRM column). Similar effects can be seen when we inspect the sensitivity of the total population size to *Q*_*ij*_ (Fig. S12B and S13B). From this analysis we conclude that we cannot expect to confidently identify all possible differentiation rates.

Next, we used simulation to investigate the practical identifiability of the differentiation pathways. We fitted model IV (CD8 lineage) and model III (CD4 lineage) to the true data to obtained parameter estimates. We then systematically perturbed the *Q*_*ij*_ for each pair of sub populations (*i, j*): using a range of 51 values between 57% and 175% of the point estimate. With each perturbed parameter value, we then generated a pseudo dataset, and fitted the model to it. We then calculated the correlation between the perturbed, ground-truth parameters *Q*_*ij*_ and the values estimated from the pseudo data (Fig. S12C and S13C). For the CD8 lineage, we clearly see that the best identifiable rates correspond to efflux from the large populations DP TRM and CD69 SP TRM, which is consistent with our sensitivity analysis. Notice that a small target population size does not preclude identifiability of differentiation rates to that population. For example, we can quite confidently say that the rate of differentiation from DP and CD69 SP TRM to DN TRM is likely small, despite DN TRM being relatively small throughout the time course. The pattern is less clear for CD4 T cells, although we mostly find a positive correlation between ground truth parameters and estimates, and the aforementioned Bcl-2hi TRM population has multiple identifiable efflux rates. Notice that we also find a number of negative correlations between the ground truth and estimates, possibly due to stochasticity.

An example of how (unperturbed) ground truth parameters correspond to estimated *Q*_*ij*_ values within a single simulated dataset is given in Fig. S12D and S13D. This shows that of the *d ×* (*d* − 1) rates, the ones that are truly large are also estimated to be large, while a clear lack of differentiation is also reflected in a small estimate.

Finally, we investigated the identifiability of the models themselves (I, II, III, IV) as opposed to the parameters. Using the four model fits to the CD8 and CD4 data (Fig. 3 and S8), we generated pseudo data using the estimated parameters. We then fit all four models to the four pseudo datasets, resulting in 16 fits. For each pseudo dataset, we then used formal model comparison to identify the best model (Fig. S12E and S13E). Most noticeably, this analysis shows that if the data is generated by model IV (the most complex), then we would select model IV with very high confidence. However, if III is the true data-generating model, then we still have a high chance of selecting model IV, especially for the CD8 lineage. Comparing this to Table S1, we see that for the CD8 lineage, we very confidently identified model IV for the real data, and give virtually no weight to model III. This pattern corresponds to row IV of the model-identifiability matrix (Fig S12E). For the CD4 lineage, we cannot very confidently choose between models II, III, and IV, which corresponds to the row III of the model-comparison matrix (Fig S12E).

#### Uncertainty in clustering highlights shortcomings of sequential approaches

Stratifying cells within the UMAP by timepoint allows one to visualize changes in the distribution of expression profiles within each cluster (Fig. 2D). Ideally, the distribution of cells within a sub-population should be independent of time, but from 20 DPI onward the distribution within CD69 SP T_RM_ shifted towards T_EM_ and CD103 SP T_RM_, T_EM_ moved towards to T_CM_, and DP T_RM_ moved away from SP and DN T_RM_. Related to these shifts, we found variation in the certainty with which cells could be assigned to each population. We quantified this uncertainty using the empirical entropy of these assignments (Fig. S3A). A low entropy corresponds to a cell that is always assigned to the same population, while high entropy indicates that different Leiden runs result in different assignments. Uncertainty of assignment also varied between populations (Fig. S3B). These discrepancies indicate that the Leiden clustering method may not capture important features of the data, and suggests that a more refined representation of the high-dimensional data is needed – one that takes into account the time evolution of the cell densities in phenotype space.

#### Integrated Approach

To deal with uncertainty in cluster assignment, we took inspiration from the single-cell modeling tools scVI [31] and scANVI [32]. We assumed that cell’s flow cytometric phenotype, *x*, can be modeled by lower-dimensional latent vector *z* representing the cell’s internal state, and that *x* can be reconstructed from *z* with a complex non-linear map, modeled with an artificial neural network (NN). The measured fluorescent intensities are noisy, and so *z* cannot be computed exactly; instead, for a given *x* we aim to generate a probability density for *z*. To avoid explicitly modeling this density for each cell, we make use of an amortized inference model, using the observations *x* to predict a parameterized distribution function for *z*. This predictor is also a NN model. In the language of variational auto-encoders (VAE), the amortization network is the ‘encoder’, and the network that generates observations *x* given a latent state *z* is the ‘decoder’ (Fig. 4A).

**Figure 4:**
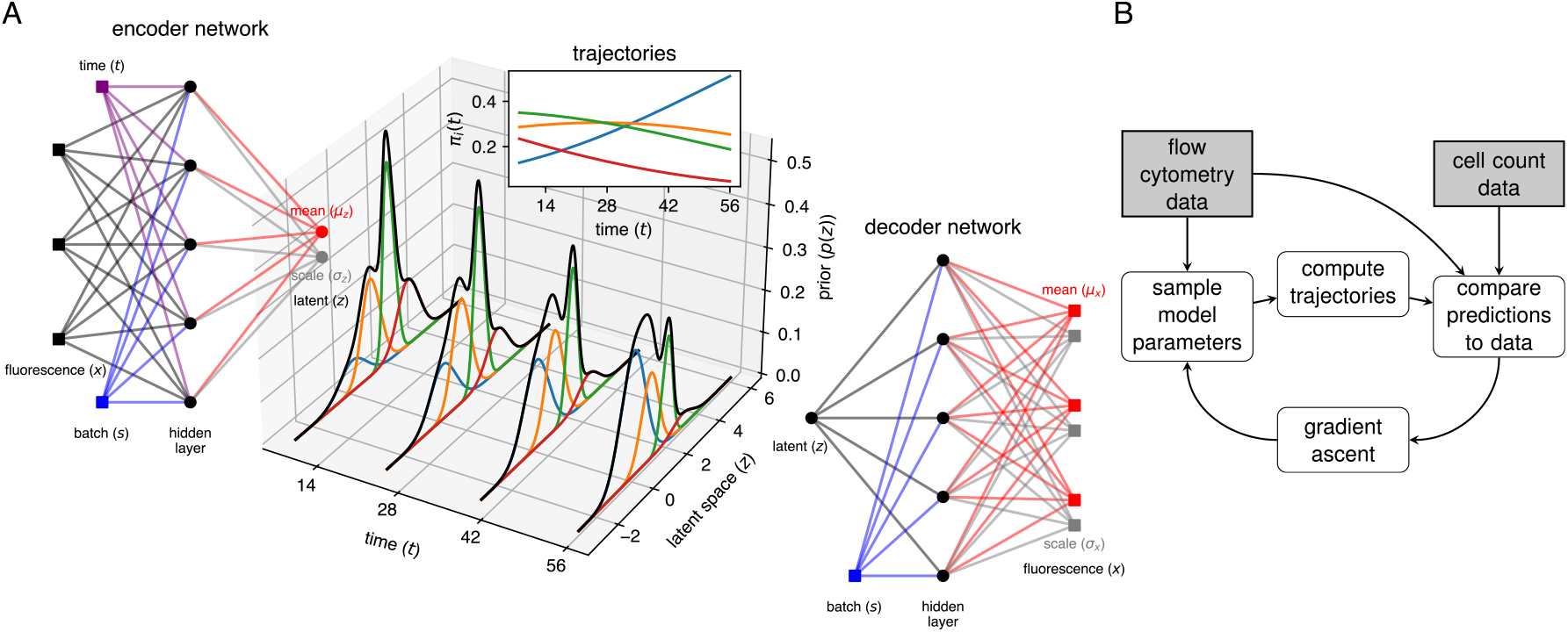
The integrated approach. **A**. Schematic of the VAE model. Using the fluorescence data *x*, we use a fully connected encoder network to infer the latent cell state (specifically, the variational parameters *µ*_*z*_ and *σ*_*z*_ for the posterior distribution of *z*|*x*; see Methods). The prior distribution of *z* is given by a Gaussian mixture model with time-dependent mixture weights determined by the dynamical model. The mixture components (multivariate normal likelihood functions) and mixture weights determine the categorical distribution of cluster membership of a cell. Using the decoder network, we compute parameters *µ*_*x*_ and *σ*_*x*_ for the distribution of *x*|*z*, thereby accounting for the measurement error in *x*. **B**. Flowchart indicating the logic behind the integrated approach. The model is fit directly to single-cell flow cytometry data and cell count data using stochastic variational inference (SVI). The posterior distribution of the model parameters (loss and differentiation rates, initial conditions, NN weights and biases, GMM parameters) are iteratively optimized. In each iteration we sample candidate parameters, and solve the system of ODEs. We then match the data with our predictions, and adjust the posterior to improve this match with the gradient ascent method.

To model the population structure within the latent space, we used a multivariate Gaussian mixture model (GMM) [33, 34]. The components and weights of the GMM correspond to the different T-cell subpopulations and their relative sizes, respectively. The ODE models predict the weights at any time point, and so the GMM describes a distribution of cell states that changes smoothly with time, parameterized by the rates of loss and differentiation, and the time-independent mixture component parameters. These mixtures overlap within the latent space, and so their time-dependent weights can capture shifting distributions of cells in both the latent and feature space. A schematic describing the time-dependent GMM is shown in Fig. 4A, where the latent space is represented by the 1-dimensional real line. A model predicting the size trajectories of four subpopulations is used to compute the weights of the mixture components (colored bell curves). The black curves show how the combined latent state distribution of all sub-populations evolves with time. The computational workflow for this method, which we refer to as the integrated approach, is illustrated in Fig. 4B. We implemented it in Pyro [35], and describe it in detail in Methods.

#### Dimension reduction using the integrated approach preserves information within the flow cytometry data

The sequential approach revealed evidence for both differentiation and time-dependent net loss rates, and so we proceeded with model IV. We first assessed our ability to find a latent representation consistent with our understanding of the marker expression data. We began with a 6-dimensional latent space representation of the 10-dimensional space of marker expression levels. Our results were similar with 5- and 7-dimensional latent spaces. Using a UMAP embedding of the latent space to visualize the learned representation, we saw that the VAE model was able to distinguish cells with different marker expression profiles (Fig. 5A) and yielded distinct populations (Fig. 5B). To visualize these sub-populations more clearly, we sampled latent vectors from each of the mixture components, mapped them the UMAP space, and calculated the contours of the 75% maximum density for each of the components. We also took the posterior samples of the location vectors of each of the components and projected them in UMAP space (indicated by colored dots in Fig. 5B). These location vectors can be interpreted as the typical cell state in a component. The contours and typical cell states match well with distinct regions in the latent space, indicating that the GMM is correctly capturing the distribution of latent cell states and, by extension, the distribution of the flow cytometry data. To account for possible batch effects, the batch information was included in the training data for the VAE. However, the trained models did not make use of large batch corrections (SI Text D), possibly due to the fact that cells from all mice were analyzed simultaneously.

**Figure 5:**
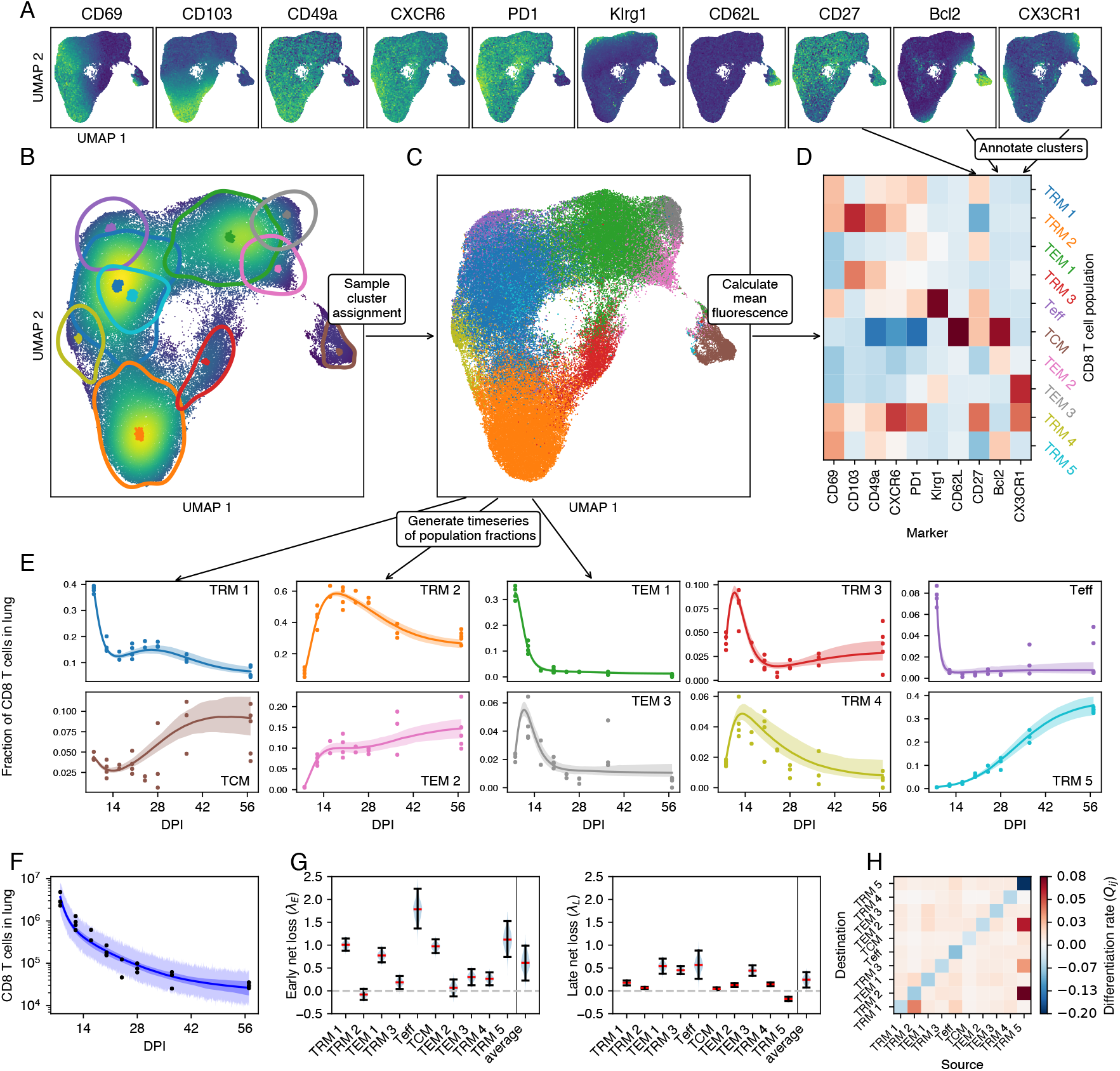
Modeling the dynamics of lung CD8 T cells with the integrated approach. Results are based on data from *n* = 27 mice. **A**. For each marker, the fluorescence intensity of each cell is displayed in a UMAP representation of the latent space. **B**. Cell density and 95% confidence ellipsoids for GMM components projected into the same UMAP space. Dots are the GMM location vectors. **C**. Using the GMM, a population is assigned to each cell. **D**. Mean fluorescence intensity of each marker and population identified by the GMM. **E**. Calculated population frequencies per mouse and timepoint, with median trajectories predicted by the model and 95% CrI (bands). **F**. Measured numbers of CD44+ CD11a+ protected CD8 T cells in lung (points), trajectory of the median prediction (line), 95% CrI (dark band), 95% posterior predictive interval (light band). **G**. Marginal posterior distributions of net loss rates during the early and late stages, with medians and 95% CrIs. **H**. Differentiation matrix.

#### The integrated approach yields finer-grained T cell phenotypes than the sequential approach

The GMM defines a distribution of cell states as a function of time, which we can match with data via the decoder NN. However, for each cell we can also use the encoder NN to compute its latent cell state, and then use the GMM to assign the cell to one of the mixture components. This is a probabilistic (or ‘soft’) cluster assignment; for each cell, the GMM gives a distribution of probabilities that the cell belongs to each component. By sampling from these distributions, we can assign to each cell a single component label (Fig. 5C). At this stage, we have effectively clustered the data, but here the key difference with the sequential approach is that this happens *after* we trained our dynamical model. The inference method does not rely at all on the fact that we assign cells to single clusters. Now that we have clustered the flow cytometry data, we can interpret the different mixture components as CD8 T cell sub-populations, and annotate them using the mean fluorescence intensities per cluster (Fig. 5D). We compared populations discovered by the two approaches using the Jaccard index (Fig. S6A). The integrated approach recovered 7 of 8 populations identified by the sequential method. Namely CD69 SP T_RM_, DP T_RM_, CD103 SP T_RM_, which we denote TRM 1, 2, 3; T_Eff_ and T_CM_; CX3CR1lo T_EM_ and CX3CR1hi T_EM_, which we denote TEM 1 and 3. It did not identify the transient DN T_RM_ subset that expressed PD-1, CXCR6, and the early activation marker CD27, and was lost quickly early in the resolution phase (Fig. 2E). Instead, the integrated approach stratified two populations further; CX3CR1lo T_EM_ were separated into Bcl-2lo and Bcl-2hi (TEM 1 and 2), and CD69 SP T_RM_ became three subsets; Bcl-2lo CX3CR1lo, Bcl-2lo CX3CR1hi, and Bcl-2hi CX3CR1lo (TRM 1, 4, and 5 respectively). This partitioning could not be readily distinguished by UMAP inspection alone (Fig. 5A).

#### With the integrated approach, data and model are entwined

Having clustered the flow cytometry data, we could again construct time series, enumerating for each mouse the number of cells per population (points in Fig. 5E), and then compare their relative sizes to the trajectories predicted by the fitted model (model IV; curves and 95% credible envelopes in Fig. 5E). This model accurately captured many features of the distribution of phenotypes, as well as the total population size (Fig. 5F). The CD69hi Bcl-2lo T_RM_ (TRM 1) declined rapidly relative to the other populations between 8 and 12 DPI, recovered, but decreased again from around a month after infection, due to the continually increasing prevalence of CD69hi Bcl-2hi T_RM_ (TRM 5). This subset was present initially at a very low frequency, but come to comprise *>* 30% of lung CD8 T cells by 2 months post infection. A similar pattern can be seen for T_EM_, which increased in frequency soon after the peak of infection but declined rapidly after 10 DPI. The Bcl-2hi T_EM_ (TEM 2) population was initially very small, but quickly replaced Bcl-2lo T_EM_ (TEM 1 and 3) and stabilized at 10% of CD8 T cells from 16 DPI. Therefore the integrated approach reveals that the Bcl-2hi and Bcl-2lo T_RM_ and T_EM_ populations, which are otherwise phenotypically similar, exhibit quite divergent dynamics. However, there is one subtlety with presenting the data as a timecourse of discrete observations. Calculating the probability that a cell is assigned to a cluster uses two pieces of information; the cell’s marker expression, and the expected sizes of each of the clusters. The latter are dependent on time, and this time dependence is in turn learned from the flow cytometry data. Therefore, the data displayed in Fig. 5E are in part generated by the model, and as a result we expect the data points to follow the model trajectories more closely than in a standard fitting approach.

#### Bcl-2-expressing T_RM_ act to sustain lung CD8 T cell populations long-term

As with the sequential approach we found that up to around 14 DPI T_Eff_ are lost most rapidly, followed by TRM 1 and T_CM_. At the same time, TRM 2 and 3, and TEM 2 and 3 were growing in numbers (Fig. 5G, left panel). From approximately 14 DPI onwards, all populations declined in number except for TRM 5, the Bcl-2hi CD69hi CD103lo subset (Fig. 5G, right panel). Bcl-2lo T_EM_ (TEM 1) were lost most rapidly, consistent with their lack of typical residency markers. Notably, at these late times T_CM_ and T_Eff_ were more persistent than canonical T_RM_. Most strikingly, Bcl-2hi CD69 SP T_RM_ (TRM 5) had a positive intrinsic net growth rate in the late memory phase (that is, we inferred that their rate of self-renewal was greater than their combined rates of death or egress from the lung), but their numbers declined due to their high rates of differentiation into DP T_RM_ (TRM 2), Bcl-2hi T_EM_ (TEM 2), T_CM_, and CD103 SP T_RM_ (TRM 3). Therefore, our analyses suggest that Bcl-2hi T_RM_ help to maintain phenotypic diversity among CD8 T cells in the lung in the weeks following IAV infection.

In our sequential analysis, we identified identifiability issues with differentiation rates. To address this in the integrated approach, we decided to leverage the latent representation learned by by the model. The idea is that distances in the latent space correspond to similarity in cell state, and that state transitions are more likely between similar states. This assumption is commonly made in other approaches to inferring differentiation pathways [36, 37]. To implement this restriction, we used a mixed-effects-type prior distribution for the differentiation rates *Q*_*ij*_ that is dependent on the distance between the mixture components of T cell populations *i* and *j*. The parameter estimate for this mixture model show that a larger distance is associated with a smaller differentiation rate (Fig. S14A and B). By inspecting the credible interval of the individual *Q*_*ij*_ as a function of the distance between component *i* and *j*, we find an inverse trend, but also that the model can account for outliers (Fig. S14C). Note that cell surface marker expression measured by flow cytometry is only a limited representation of the cell state.

#### Model validation with the integrated approach

In the integrated approach the generated and predicted cluster frequencies are interdependent to an extent. How then do we validate a model’s utility or predictive capacity? In contrast to the sequential method, which predicts only the trajectories of the sizes of predefined clusters, the VAE model is generative – that is, it can be used to predict the trajectory of the flow cytometry data itself. Such a trajectory is shown in Fig. 6A. Each square shows a UMAP of the latent space at a particular time post infection. The squares highlighted in grey are UMAPs of the actual data, and the others are simulated by the model. This series of UMAPs acts as a visual posterior predictive check (i.e. a ‘model fit’), analogous to a visual assessment of how well the curves in Fig. 3B capture the trends in the datapoints. In this case we can see the method describes the data well, and highlights time periods that may require denser sampling. For instance, the simulated data between 8 DPI and 12 DPI show rapid shifts in the distribution of phenotypes; CD69 SP T_RM_ rapidly disappear, DP T_RM_ increase in frequency, and then T_EM_ start to decline. This order of events is not directly observed, and might be resolved with intermediate samples.

**Figure 6:**
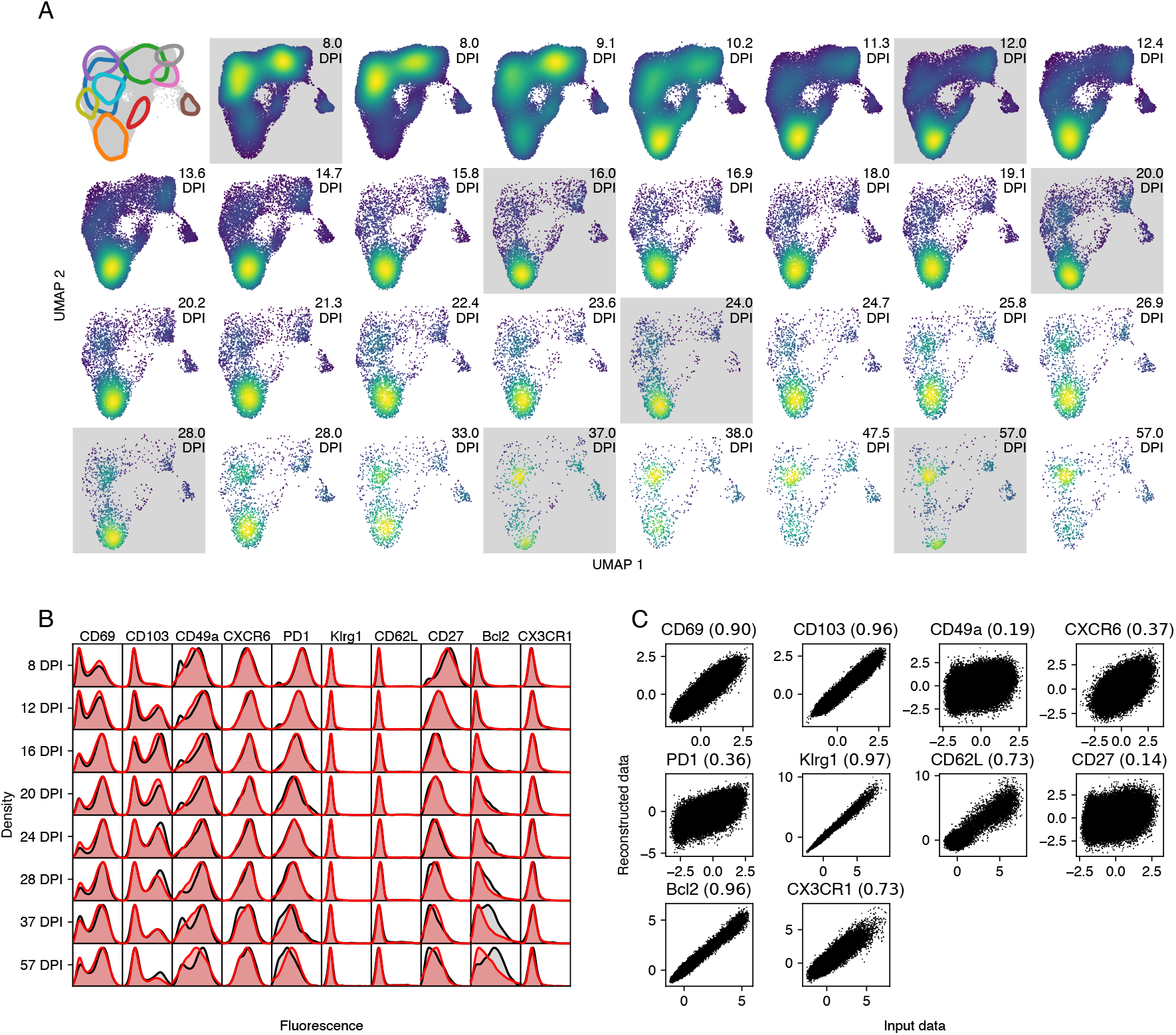
Posterior predictive checks using simulated marker expression data. Results are based on data from *n* = 27 mice. **A**. UMAPs of marker expression data, either simulated using the fitted model (white background), or representing the observations (grey background). The location of the inferred clusters are highlighted in the top left panel (colors as in Figure 5B). The color (blue to yellow) shows the density of the points in UMAP space. The number of sampled cells is gradually decreased with DPI to match the sample size of the actual data. The marker expression data (simulated or real) is first mapped to the latent space using the trained VAE model. Then, a UMAP model is trained on the combined true data. The same UMAP model is used to map the simulated data to the plane. **B**. Marginal distributions of marker expression (cf. Fig. 1H). Data is shown in black, simulated data is shown in red. **C**. Input data and reconstruction using the auto-encoder model. The number in brackets is the coefficient of determination (*R*^2^).

Because the integrated approach models the joint density of the marker expression, we can easily marginalize to single cell surface markers and compare the model predictions with the data. This provides another posterior predictive check (Fig. 6B). Pseudo-observations are created by sampling from the GMM at the observation times, and using the decoder network to transform the latent space to the feature space. The marginal distributions of the pseudo observations (red density plots) largely coincide with the true data (black densities), but there are some discrepancies. Most notably, the VAE has difficulties capturing the bimodal shape of the CD49a density, as well as the shape of the Bcl-2 distribution at late time points.

To investigate these discrepancies further, we made use of the auto-encoder architecture of the model. An auto-encoder finds a low-dimensional representation of the data, thereby removing noise and finding essential features [38]. We can test how well our auto-encoder has learned to represent the data by first encoding the marker expression data set, and then reconstructing the expression profiles from the latent cell states using the decoder network. For each marker, we plot the observed fluorescence intensities against the reconstructed values (Fig. 6C). A perfect reconstruction would yield a perfect correlation between input and output. For the markers CD69 and CD103, we find a very high correlation (*R*^2^ = 0.90 and 0.96, respectively), while for CD49a and CD27, we find low correlations (*R*^2^ = 0.19 and 0.14, respectively). Hence, it appears that the VAE discarded most of the information encoded by CD49a and CD27 expression, indicating that variation in these markers does not relate strongly to heterogeneity in the dynamics of CD8 T cell subsets in the lung.

#### The dynamics of lung-localized CD4 T cells following IAV infection

We then applied the two approaches to the CD4 T cell data. With Leiden clustering and consensus annotation, we identified eleven distinct populations (Figs. S7A and B). Seven clusters possessed a T_RM_ phenotype, delineated by varying expression levels of canonical T_RM_ markers CD69, CD49a, CXCR6, CD103. One of the T_RM_ subsets also expressed KLRG1, suggesting an effector phenotype. We also identified four populations lacking these residency markers: a CD62Lhi TCM-like subset, a subset expressing high levels of FR4 and PD-1, characteristics of T follicular helper (T_FH_) cells in the lung [39], and two populations with low expression of all markers measured, distinguished only by the presence of Bcl-2, which we termed T_EM_.

As we found for the CD8 T cells, the joint distribution of marker expression on CD4 T cells gradually shifted with time post infection (Fig. S7C). The peak of the response was dominated by Bcl-2lo T_EM_ and the CD69lo CD49alo T_RM_ population, which expressed high levels of PD-1 and CD27, markers upregulated during activation (Fig. S7D). By 3 weeks post infection, CD69hi CD49alo and CD69hi CD49ahi T_RM_ dominated, giving way in turn to two Bcl-2-expressing T_EM_ and T_RM_ subsets by 2 months post infection.

We fitted our four model variants to the timecourses of total CD4 T cell numbers (Fig. 1C) and the subset frequencies (Fig. S7D). Visual inspection and posterior predictive checks indicated that all four models fit the data reasonably well (Fig. S8A), but formal model comparisons indicated that time-dependent net loss alone (model II) or differentiation alone (model III) provided sufficient descriptions of the data (Table S1).

As was the case for CD8 T cells, all models exposed remarkable heterogeneity in the dynamics of CD4 T cell subsets in the lung after the resolution of IAV infection (Fig. S8B; using model II), particularly among the T_RM_-like populations. The loss rates estimated with model III showed a similar, but slightly more homogeneous pattern (Fig. S8C). Again using model III, we found evidence for ongoing differentiation from CD69lo CD49alo T_RM_ to CD69hi CD49ahi T_RM_ and CD69lo CD49ahi T_RM_, and from Bcl-2hi T_RM_ to multiple T_EM_ populations and effector cells (Fig. S8D). Finally, the data suggests a pathway from CD69hi CD49alo T_RM_ to T_EM_. These data indicate the dynamically shifting T cell populations within the local environment, even after resolution of infection. High entropy scores within T_RM_ clusters indicated substantial uncertainty in Leiden cluster assignment for CD4 T cells (Fig. S3D-E), underscoring the need for integrated analysis to dissect the subtle changes in subset composition.

Next, we used the integrated approach to refine the model fit for the CD4 data. We chose to continue with model III as it performed just as well as the full model and enabled us to further explore differentiation pathways. The VAE generated a latent representation that respected marker expression, and the location of the mixture components in UMAP space corresponded well with high density regions (Fig.7A-B). Using mean fluorescence intensity (Fig. 7C), the integrated approach identified T_CM_ (CD62Lhi) and a KLRG1hi subset that also expressed T_RM_ markers (TRM 7) whose phenotypes were highly concordant with those identified using the sequential approach. In contrast, many of the T_RM_ and T_EM_ populations were distinct between the two approaches (Fig. S6B).

**Figure 7:**
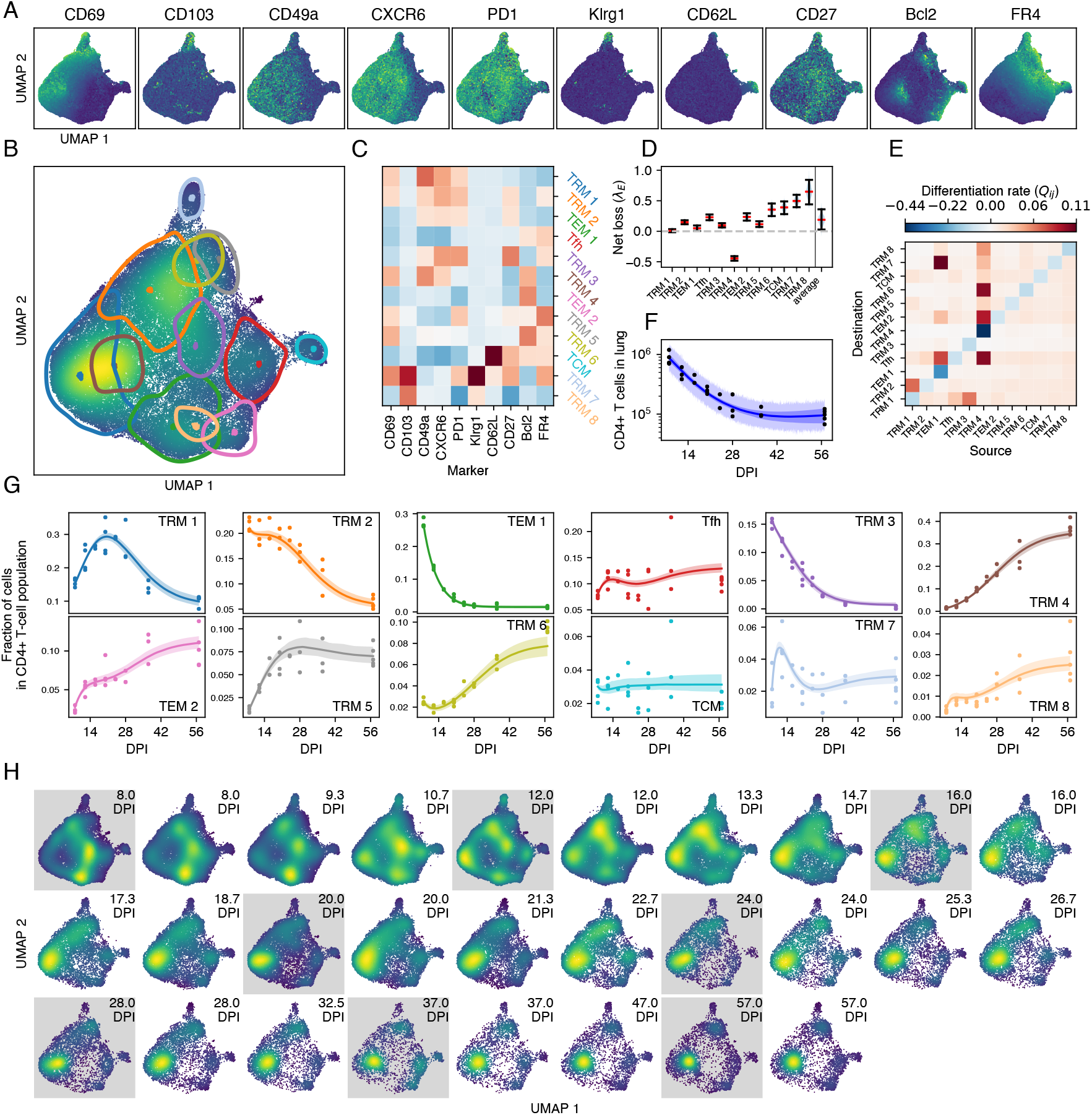
Modeling the dynamics of lung CD4 cells with the integrated approach. Results are based on data from *n* = 27 mice. **A**. For each marker, the fluorescence intensity of each cell is displayed in a UMAP representation of the latent space. **B**. Cell density and 95% confidence ellipsoids for GMM components projected into the same UMAP space. Dots are the GMM location vectors. **C**. Mean fluorescence intensity of each marker and population identified by the GMM. **D**. Marginal posterior distributions of net loss rates, with medians and 95% CrIs. **E**. Differentiation matrix. **F**. Measured numbers of CD44+ CD11a+ protected CD4+ T cells in lung (points), trajectory of the median prediction (line), 95% CrI (dark band), 95% posterior predictive interval (light band). **G**. Calculated population frequencies per mouse and timepoint, with median trajectories predicted by the model and 95% CrI (bands). **H**. UMAPs of marker expression data, either simulated using the fitted model (white background), or representing the observations (gray background).

The integrated approach identified eight T_RM_ clusters, defined by varying combinations of T_RM_ markers. In contrast to the sequential approach, several of these clusters (TRM 2, 3, 5, 6 and 7) expressed the T_FH_ marker FR4, which was restricted to T_FH_, T_EM_, and T_CM_ in the sequential approach. The remaining T_RM_ clusters (TRM 1 and 4), were presumably canonical T_h_1-like cells characteristic of a typical IAV response. Again, these subpopulations exhibited diverse loss rates (Fig. 7D), with the KLRG1-expressing TRM 7 and 8 lost at the highest rate, and TRM 4 (characterized by CD69, CD49a and Bcl-2) showing an intrinsic net positive growth rate.

We found evidence for more flows from FR4lo to FR4hi subsets than *vice versa*, indicating fluxes towards populations with characteristics of T follicular helper cells (TEM 1 to TRM 7 and T_FH_; TRM 1 to TRM 2, TRM 4 to T_FH_ and TRM 6). However, these populations had relatively high loss rates and as the population declined as a whole (Fig. 7F) they were outcompeted by Bcl-2-expressing subsets (TRM 4, and TEM 2; Fig. 7G), akin to the long-lived T_RM_ in the CD8 data. In contrast to the CD8 lineage, we did not find a relation between distance in the latent space and differentiation rates (Fig. S14B and D), likely reflecting the tighter clustering of CD4 T cell phenotypes.

As described for CD8 T cells, the generative nature of the VAE provided a visual predictive check (Fig. 7H), showing an excellent correspondence between the data and the fitted model. The observed and predicted marginal distributions for surface markers corresponded well (Fig. S9A), with the exception of CD49a and CD27. These discrepancies were also noticeable when we reconstructed data using the VAE (Fig. S9B) where CD49a and CD27 had the lowest coefficient of determination (*R*^2^ = 0.16 and 0.12, respectively). In contrast, CD69, Bcl-2 and FR4 were almost perfectly reconstructed (*R*^2^ *>* 0.97).

## Discussion

Developing methods to connect time series of high dimensional data with mechanistic, dynamic models of cell behavior is a pressing challenge. Here we described a straightforward ‘sequential’ approach in which one clusters and then models the occupancy of these clusters over time. Intuitively, though, this approach may be problematic when phenotypic diversity within a population does not permit clear separation of discrete subpopulations. To address this issue we explored a more holistic approach which allows for timevarying uncertainty in the assignment of cells to clusters, simultaneously modeling the cellular dynamics and its low dimensional representation.

In the setting of the T cell response to influenza A virus infection, the sequential approach showed that the net loss rates of CD8 T cells within the lung are strongly dependent on their phenotype and on time, and also found evidence for ongoing differentiation between cell types. The integrated approach partitioned CD8 T cell phenotypes and their dynamics in a similar way, but also provided more resolution. Specifically, we found that lung-resident CD8 T cells appear to be maintained by a Bcl-2hi CD69hi subset that lacks expression of the canonical T_RM_ marker CD103. Expression of Bcl-2 may be important for counteracting pro-apoptotic signals arising from the nutrient-poor environment of the airway[40]. It has been suggested that CD69hi CD103hi T_RM_ in the airway are maintained by T_RM_ in the lung parenchyma and that migration is mediated by increased CXCR6 expression[12]. Although here we did not perform bronchoalveolar lavage (BAL) to study cells within the airway directly, we did find evidence for differentiation within the parenchyma from these Bcl-2hi CD69hi T_RM_ into CD69hi CD103hi (DP) T_RM_, which also co-expressed CXCR6 at intermediate levels.

We found that CD4 T cells within the lung were also dynamically and phenotypically heterogeneous, but with subsets less well defined than their CD8 T cell counterparts. In contrast with the CD8 results, the CD4 T cell data could be explained by either time-dependent loss of independent populations, or constant loss of differentiating populations. As with CD8 T cells, however, we found that Bcl-2-expressing populations come to dominate the CD4 T cells within the lung at later time points, and that these cells might be a source for other populations, thereby sustaining heterogeneity. Notably, we also saw evidence for one-way differentiation from canonical T_RM_, which were largely T_h_1-like, to FR4-expressing cells that were more T_FH_-like.

Our flow cytometry panel contained five markers commonly associated with tissue residency (CD69, CD103, CD49a, CXCR6, and PD1), and others that are typically not expressed on canonical T_RM_ (CD62L, CX3CR1, and KLRG1). T_RM_ have been sub-classified further with markers such as CD101, CXCR3, and CXCR4 [14], and several others [41]. Expanding the panel will likely reveal more fine-grained subpopulations, each with distinct dynamics. For instance, it has been shown that T_RM_ in the airway rapidly lose their CD11a expression [12]. Airway T_RM_ also show distinct dynamics as they are lost more rapidly than T_RM_ in the parenchyma [40]. Here, we used CD11a to pre-select antigen-experienced cells in our gating strategy. If data from BAL were included, adding CD11a to the training data instead could be beneficial. Our inferences could also benefit from inclusion of the Ki67 marker, which is expressed transiently after cell division. Using this marker could allow us to disentangle net loss rates (*λ*) into division (*ρ*; Fig. 3A) and death or egress (*δ*). However, this would require more complex models, and will be left for future work.

A typical workflow may benefit from using both the sequential and integrated methods. The sequential approach makes it easy to visualize the data as a set of timeseries, and hence aids in the the model development phase, and may be sufficient if the subsets are very clearly defined (such as the T_CM_ populations we identified here). The integrated approach is particularly powerful when standard clustering approaches do not clearly delineate phenotypes. In any case it is well suited for fine tuning of the model, parameter estimates and the population structure. Here we used the sequential approach for model selection and explored the integrated approach only with the favored models for CD8 and CD4 T cell dynamics. As the integrated approach is likelihood-based it should also lend itself to model comparison, but performing this is difficult for variational methods and remains an open challenge[42]. Facilitating model selection within the integrated approach would provide significant advantages. In the sequential approach, it is not possible to apply standard selection criteria to establish the optimal number of clusters, because the clustering step occurs before model fitting, and so different numbers of clusters represent entirely distinct datasets. In contrast, the integrated approach could provide an objective measure of the number of subpopulations present in the data, because it deals with the likelihood of the observations themselves.

Increasing the number of clusters resolved new and distinct populations, such as a CX3CR1hi effector-like subset, and a Bcl-2hi subset of the DP T_RM_; However it also generated in very small and likely spurious populations that were present at single time points, possibly due to batch effects. Rational determination of the optimal number of clusters is therefore an important future research direction.

A related issue with the integrated method is that, because models are fitted to high-dimensional data, it is challenging to perform posterior predictive checks. We used two-dimensional UMAP projections to address this issue (Fig. 6A), comparing the density of the observations in UMAP space with the density of pseudo-observations generated by the model at intermediate time points. A recent analysis of dimension reduction methods argued that for high-dimensional single-cell data the density in UMAP space corresponds poorly with cell density in the feature space [43]. However, the dimensionality of our data is an order of magnitude lower than the data used in their analyses. Therefore we are confident that, in the realm we explored here, using UMAP for posterior predictive checks is a reasonable procedure.

In both approaches, the model equations take the same functional form for all populations. In general this constraint will be too stringent. For instance, during the expansion phase of the immune response diversifying subsets of T cells may exhibit highly distinct dynamics, which may need to be modeled with structurally different equations. Further, programmed bursts of division characteristic of clonal expansion phase require multiple equations to describe a single population, one for each round of division [44, 45]. Structural heterogeneity within models is easily accommodated by sequential approaches, because one assigns phenotypes to each of the clusters before model fitting. It is more problematic for the integrated approach; because we do not know *a priori* which cells will end up in which mixture component, an equation may end up describing the wrong cluster of cells. A potential solution is to use semi-supervised clustering [32]. Here we assign to a small set of cells a phenotype that we want to model with a particular type of equation. We then add a term to the likelihood function that makes it much more likely that those selected cells end up in the appropriate cluster.

Our approach is similar to scVI [31] and scANVI [32] in its use of a VAE, although our method is currently designed for flow cytometry data instead of very high dimensional single-cell RNA sequence data, which requires more complex error models. The use of hierarchical Gaussian mixture models for the analysis of flow cytometry data has been explored elsewhere [46], but here we fit a GMM on a lower-dimensional state space after applying a non-linear transformation. The transformed data might be more easily described by a GMM than the raw flow cytemetry data, especially when the latter is high-dimensional. Autoencoders for single-cell data analysis were first introduced as a de-noising and imputation tool[47], but have evolved rapidy. Currently there exists a large collection of methods based on variational autoencoders [31] which can analyze scRNA-seq data and other modalities. These methods can roughly be grouped into ‘exploratory’ and ‘guided’ [48]. Exploratory methods do not use any covariates of interest to guide latent state recovery or community detection, while guided methods explicitly incorporate such data [49]. Our integrated method falls into the guided class of models, as we use time information directly in the data manifold reconstruction, as well as community detection steps.

For the combination of models and data we studied here, parameter identifiability was a legitimate concern. We are trying to infer both the differentiation rates between populations and their net loss rates, from time series of their sizes. Although we showed that with sufficient data these quantities can be distinguished, in practice not all are equally identifiable. In particular, it can be difficult to reliably quantify the flows from small T cell populations into larger ones. The T_RM_ differentiation pathways identified by this study should therefore be verified with further experiments. One could also put constraints on possible pathways with additional data from fate-mapping mouse models [50, 51] or with developmental trajectories inferred from CITE-seq data [37, 52].

Understanding the dynamical properties of immune responses is key to solving many fundamental problems, ranging from finding cures to chronic diseases such HIV-1 infection, designing vaccines that provide heterosubtypic immunity to respiratory viruses such as influenza, and immuno-therapies targeting tumors. Continuing technological advances are yielding bountiful data at the single cell level, resolving phenotypes at unprecedented scales and promising to advance our abilities to tackle these problems. Meanwhile, the deep learning revolution is presenting us with an expanding suite of tools to process and visualize and interpret these data. Mathematical modeling has a long tradition of aiding the interpretation of immunological and virological data, but has largely relied on heavily curated datasets. The approaches we present in this paper, and others like them, bring dynamical models closer to the observations. This bridging will increase the utility of mathematical models for shaping and testing our intuitions regarding biological processes.

## Methods

### Experimental procedures

#### Mice

Mice were housed in specific pathogen–free conditions in the animal facilities at Columbia University Irving Medical Center (CUIMC). Eight week old female C57BL/6 were purchased from the Jackson Laboratory. Infections were performed in biosafety level 2 biocontainment animal facilities. All animal studies were approved by Columbia University Institutional Animal Care and Use Committee.

#### Influenza infection

C57BL/6 mice were briefly anesthetized using inhaled isoflurane and infected intranasally with X31 influenza A virus at a 50% tissue-culture infectious dose (TCID50) of 5000 in 30μl of PBS. Infected mice were weighed daily and animals that lost more than 30% of their starting weight were humanely euthanized. The infections were staggered such that all samples were processed and analyzed on the same day, to mitigate batch effects.

#### Tissue preparation

Mice were intravenously injected with 5μg of Brilliant Violet (BV) 421-conjugated anti-CD45.2 antibody 5 minute before cervical dislocation. Spleen and mediastinal lymph nodes (MedLNs) were processed by mechanical disruption and passed through 70μm filters (Corning). Singlecell suspensions of lungs were prepared first be mechanical disruption followed by digestion with 1mg/ml collagenase and DNAse (Sigma) for 40 minutes at 37°C. Lungs were then passed through a 70μm filter. Red blood cells were lysed from the spleen and lungs using ACK lysis buffer (Gibco).

#### Flow cytometry

Single cell suspensions were washed in PBS before staining with Zombie NIR viability dye (1/1000 in 100μl PBS) for 30 minutes at 4°C. Cells were washed twice and stained with 0.5μl of PE-labeled IAb/NP_311-325_ (CD4) and 1μl of APC-labeled H-2Db/NP_366-374_ (CD8) tetramers (National Institute of Health (NIH) Tetramer Core Facility (NTCF)) in 50μl FACS buffer (PBS with 2% heat inactivated FBS + 0.05mM EDTA). For surface staining, fluorochrome-conjugated antibodies (STAR Methods) in FACS buffer were added to cell suspensions for 20 minutes at 4°C protected from light, followed by washing with FACS-buffer. Cells were then fixed and permeabilized with the FoxP3 fix/perm kit (Tonbo) for 30 minutes at 4°C. Following fixation, cells were washed with FACS buffer, and stained with antibodies for intracellular detection (STAR Methods) in Perm Buffer (Tonbo) for 20 minutes at 4°C, followed by washing and resuspension in FACS buffer. Stained cells were acquired using the Cytek Aurora flow cytometer and analyzed using FlowJo version 10 software (Tree Star).

#### Congenic transfer experiment

Three CD90.1 and twelve CD90.2 6-8 week old C57BL/6 male mice were infected with X31 influenza virus (TCID50 5000 in 30 μl). At day 14 of infection, mediastinal lymph nodes and spleen cells were pooled from the three CD90.1 infected mice. From these two groups of pooled cells, T cells were isolated using a Stemcell Technologies EasySepTM T cell isolation kit. 10^6^ pooled T cells were transferred into the infected CD90.2 mice via intraperitoneal injection and tissue was analyzed at days 2, 4, and 6 post-transfer (*n* = 3 per time point). As a control, 10^6^ pooled T cells from age- and cage-matched uninfected CD90.1 mice were transferred into D14 infected CD90.2 mice and tissue was isolated at 2 days post-transfer (*n* = 3).

### Models

#### Sequential approach

The classical analysis pipeline for our data is as follows. First, the data is pooled and clustered using a heuristic clustering method. Second, for each mouse, the number of cells in each of the clusters is counted. As the samples were taken at different times post infection, this results in a time series of cell counts per cluster. Third, dynamical models are fitted to these time series. The resulting parameter estimates and model comparison can give clues about the dynamics of the different T-cell populations in the lung.

For the first step (clustering), we used the established Phenograph method [53], which first constructs a *K* nearest-neighbor graph, and then uses the Leiden algorithm [22] for community detection. We ran the algorithm with number of neighbors set to 10 and resolution parameter set to 0.8. We used the implementation from the scanpy package [54].

For the third step (fitting models), we used the probabilistic programming language Stan (cmdstan version 2.34) [23, 24]. Bayesian inference was done with the NUTS algorithm, using 5 parallel chains, 500 warm-up iterations and 500 posterior samples. To prevent divergent transitions, we set the “adapt delta” parameter to 0.9.

#### ODE models

We developed a number of different models, each making different assumptions about T-cell dynamics and differentiation. The most general model is given in terms of the following system of ODEs

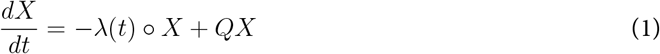

and initial condition *X*(*t*_0_) = *X*_0_. Here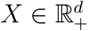 is a vector of population sizes for each of the *d* clusters. The vector *λ* contains the net loss rates for each of the clusters, and ° denotes the Hadamard product (element-wise multiplication). The matrix *Q* determines T-cell differentiation, and *Q*_*ij*_ is the per-capita rate at which cells in cluster *j* differentiate into cells in cluster *i*. The matrix *Q* is a stochastic kernel in the sense that the columns add up to zero. Hence, the diagonal elements *Q*_*ii*_ ≤ 0 denote the total efflux from cluster *i* due to differentiation.

We consider two different forms of the net loss rate *λ*. The simplest form is time-independent loss *λ*(*t*) = *λ*_*E*_. This means that the loss rates of each population stays constant over time. However, each population can have its own distinct loss rate. For instance, effector cell counts might decrease faster than T_RM_ or T_CM_ cells.

In addition to a time-homogeneous (or autonomous) model, we also consider a model in which the net loss rate decreases exponentially to a smaller loss rate.

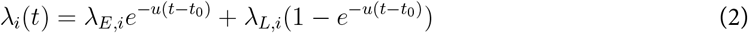

The indices *E* and *L* refer to ‘early’ and ‘late’ stage dynamics. The speed at which this switch happens (*u*) is shared between clusters. In both the homogeneous case and the inhomogeneous case, we can compare models in which we set *Q* = 0 (i.e. the clusters form independent populations) to models in which we estimate the off-diagonal coefficients of the matrix *Q* (i.e. allowing for differentiation between clusters).

In the special case when either *Q* = 0 or *λ* does not depend on time, the initial value problem (IVP) given by Eqn. (1) has a simple closed form solution. This is convenient for inference, as we will not have to use numerical integrators (and in particular sensitivity or adjoint equations). The solution is given by

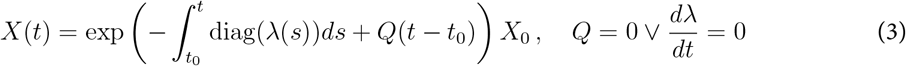

where exp denotes the matrix exponential, and diag(*λ*) denotes a square, diagonal matrix with vector *λ* on the main diagonal. The integral of *λ* is given by

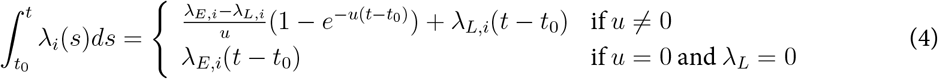

In general, we have to resort to numerical integration to solve IVP (1). To integrate ODEs in a manner compatible with the Pytorch and Pyro frameworks, we use the torchode package [55]. Specifically, we use the Dormand–Prince method with the adjoint method for back-propagation. For the sequential approach we use the built-in Dormand-Prince ODE solver in Stan.

#### Observation (or error) model and prior distributions

In order to compare our dynamical model to the data, we need to specify a likelihood function. We have two data streams: For each sample (mouse) *s*, we have the number of cells per cluster *i*, denoted *K*_*s,i*_. Additionally, we have an estimate of the total number of T cells in the tissue, denoted *M*_*s*_. According to the model, the true cluster frequencies are given by *π*_*i*_(*t*_*s*_) = *X*_*i*_(*t*_*s*_)*/Y* (*t*_*s*_), where 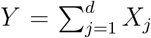 denotes the total number of T cells. We then assume that the the count vector *K*_*s*_ is sampled from a Dirichlet-multinomial distribution

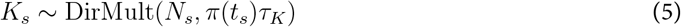

where 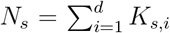 is the total number of T cells in sample *s*, and *τ*_*K*_ is a dispersion parameter. Explicitly, the probability of sampling *K*_*s*_ is equal to *N*_*s*_*B*(*N*_*s*_, *τ*_*K*_)*/ ∏ i*:*Ks,i>*0 *K*_*s,i*_*B*(*K*_*s,i*_, *τ*_*K*_*π*_*i*_(*t*_*s*_)), where *B* is the Beta function. The dispersion parameter *τ*_*K*_ determines the over-dispersion of the Dirichletmultinomial distribution, which converges to a multiniomial distribution as *τ*_*K*_ → ∞. We use an over-dispersed distribution because of the significant biological and technical variation between our samples [56].

We assume that the total number of cells in the tissue (*M*_*s*_) has a log-normal distribution

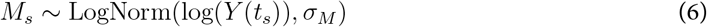

where *Y* is the total number of cells predicted by the dynamical model. The scale parameter *σ*_*M*_ is determined by the technical and biological variation between the experiments.

In addition to a likelihood function, we have to specify prior distributions for the parameters, which are listed in Table S3. As we do not force the net loss rates *λ*_*E*_ and *λ*_*L*_ to be positive, the model space includes non-biological instances in which the T-cell population grows exponentially at long time scales. To discourage this, we add a penalty term to the prior distribution. The long-term dynamics is linear and governed by the matrix *A* = *Q* − diag(*λ*_∞_), where *λ*_∞_ = lim_*t*→∞_ *λ*(*t*). The linear system *dX/dt* = *AX* is stable if the spectral abscissa *α*(*A*) ≡ max_*i*=1,…,*d*_{(*χ*_*i*_)} is negative, where *χ*_*i*_ are the eigenvalues of *A*. We therefore add a term − max{0, 10*α*(*A*)}2 to the log-prior distribution to discourage unbounded growth.

#### Model comparison

For model comparison, we used the Pareto-smoothed importance sampling (PSIS) approximation of leave-one-out (LOO) cross validation [28]. We used the method implemented in the python package Arviz [57]. The PSIS approximation breaks down for highly influential observations, and Arviz package provides a diagnostic for when this happens. For these observations, we replaced the PSIS approximation with the true LOO-CV statistic by re-fitting the model.

#### Parameter identifiability

We assessed structural identifiability with the Julia package `StructuralIdentifiability.jl` [58]. To use this package, we first have to cast the system (1) into an autonomous system of ODEs. We observe that 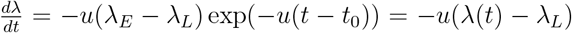 and hence the extended IVP

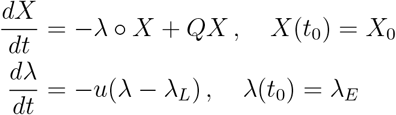

is equivalent to (1). We checked that for *d* = 2, 3, …, 12 all parameters (i.e. *λ*_*E*_, *λ*_*L*_, *X*_0_, *u, Q*) are globally identifiable.

#### Integrated approach

We estimate parameters of our ODE model and a set of “nuisance” parameters (NN weights and biases, GMM location vectors) simultaneously in a variational inference framework. We approximate the joint posterior distribution of the latent cell states, the ODE model parameters, and the GMM components with a so-called variational distribution, and optimize the variational parameters using stochastic gradient ascent. This procedure is outlined in Fig. 4B. The flow cytometry data is used together with the encoder network to generate a random sample of latent cell states *z*. At the same time, we also sample ODE parameters and GMM location parameters from the variational distribution. Using these parameters, we predict the trajectories of the T-cell population sizes at each sampling time. The total predicted population size is used to compute the likelihood of the total cell counts. The relative population sizes give us the weights of the GMM model, and allow us to compute the prior density for each latent cell state. The decoder network is used compute the likelihood of the flow cytometry data, given the sampled cell states. The prior density and likelihood are combined into the evidence lower bound (ELBO), the objective function of variational inference, which we explicitly define below. Using the gradient of the objective function, the variational parameters are perturbed in the direction of a larger ELBO at which point we can sample new parameters. We repeat this procedure until the ELBO converges.

In our integrated approach, we combine clustering with fitting of the dynamical model in a single machine learning algorithm. We use a variational auto-encoder (VAE) to infer a lower dimensional representation *z* of the marker expression data *x*. We then simultaneously fit a Gaussian mixture model (GMM) to the latent vectors *z* [33]. Each of the *d* components of this GMM represents a T-cell cluster. This means that for each data point *x*_*n*_, we can compute the probabilities *p*_*n,i*_ that cell *n* belongs to population *i* using Bayes rule

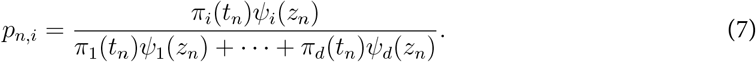

In this equation, *ψ*_*i*_ denote the probability density functions of the Gaussian mixture components, each parameterized with a mean vector *µ*_*i*_ and covariance matrix Σ_*i*_, and *π*_*i*_ represents the weight of mixture component *i*. In our method, *π* is a function of time and is determined by a dynamical model as in the sequential approach: *π*(*t*) = *X*(*t*)*/Y* (*t*).

The VAE consists of a decoder network *D* and a encoder network *E*. The decoder network *D* : (*z, s*) 1→ (*µ*_*x*_, *σ*_*x*_) takes as an argument a latent vector *z* and the one-hot-encoded batch information *s*, and returns vectors *µ*_*x*_ ∈ ℝ*m* and 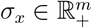 which we can use to sample a vector *x* in the feature space.

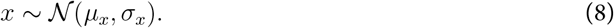

or compute the likelihood of an observed expression vector *x*, given latent vector *z*. The decoder network *D* has one hidden layer with 20 nodes and a softplus non-linearity. This hidden layer is shared by the two output layers for *µ*_*x*_ and *σ*_*x*_. We use a softplus link function to ensure that the scale *σ*_*x*_ is positive. The encoder network *E* : (*x, s, t*) 1→ (*µ*_*z*_, *σ*_*z*_) is similar in architecture to *D*, but takes as an input the expression vector *x*, the batch information *s*, and the sampling time *t* and returns a location and scale vector for the latent vector *z*, distributed as *z* ~ 𝒩 (*µ*_*z*_, *σ*_*z*_). The encoder depends on the sampling time, because it approximates the posterior distribution of *z*, which in turn depends on the time-dependent prior distribution (the GMM).

#### Target of optimization: ELBO

The goal of variational inference is to find an approximate posterior distribution *q* that is as close as possible to the true posterior, *p*. The Kullback-Leibler (KL) divergence provides a measure of their similarity;

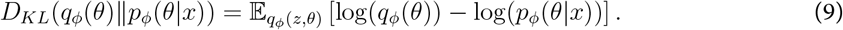

The posterior distribution is given by

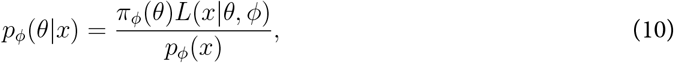

where *p*_*ϕ*_(*x*) is the evidence of the model, given the data. We would like to find a parameter vector *ϕ* that maximizes this evidence, but unfortunately *p*_*ϕ*_(*x*) = 𝔼_*π*_*ϕ*(*θ*)[*L*(*x*|*θ, ϕ*)] is generally intractable and difficult to approximate directly. This is where the variational distribution is useful. As the KL-divergence is positive, we have

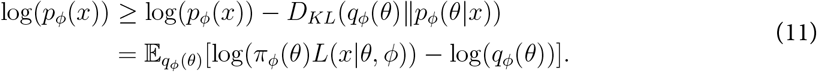

This final quantity is called the evidence lower bound (ELBO). If we then maximize the ELBO, we both minimize the KL-divergence, and given that our variational distribution is close to the posterior, we also maximize the (log) evidence of the model.

To specify the ELBO for our model we have to define three quantities: the prior *π*_*ϕ*_, the likelihood function *L* and the variational distribution *q*_*ϕ*_. First, the prior factors into three components

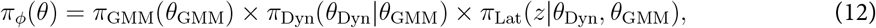

corresponding to the GMM parameters, the prior for the dynamical model parameters, and the prior for the latent vectors *z*, respectively. The parameters for the GMM consist of the location (or mean) vectors *µ*_*i*_ in the latent space of the Gaussian components, and their covariance matrices Σ_*i*_. The component weights are not free parameters, but are derived from the dynamical model. The location vectors have prior distributions *µ*_*i*_ ~ *𝒩*_*k*_(0, *I*_*k*_), where *I*_*k*_ is the *k*-dimensional identity matrix. To reduce identifiability issues, all components *i* have covariance matrix Σ_*i*_ = *I*_*k*_. Furthermore, let *D*_*ij*_ = *‖ µ*_*i*_ − *µ*_*j*_ *‖* _2_ denote the Euclidean distance between location vectors *µ*_*i*_ and *µ*_*j*_. To avoid overlap of any two mixture components (*µ*_*i*_ ≈ *µ*_*j*_), we add the penalty term 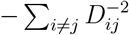to the log-prior probability.

The priors and parameters *θ*_Dyn_ for the dynamical model are specified in Table S3. To restrict the space of possible differentiation pathways in an unsupervised manner, we considered a model in which the prior probability of the differentiation rate *Q*_*ij*_ is dependent on the similarity between clusters *i* and *j*. This assumes that cell is most likely to differentiate into a state similar to its original state [36, 37]. As a measure of similarity we use the Euclidean distance between the component means in the latent space *D*_*ij*_ = *‖ µ*_*i*_−*µ*_*j*_ *‖*_2_. We take as prior distribution for the differentiation matrix 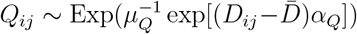 where 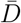 is mean all distances, and *α*_*Q*_ *>* 0 is an estimated effect of distance on the differentiation rate. Notice that with this parametrization a large distance *D*_*ij*_ implies that the differentiation rate *Q*_*ij*_ is *a priori* small.

Finally, the prior for the latent vectors *z* is given by the GMM with density

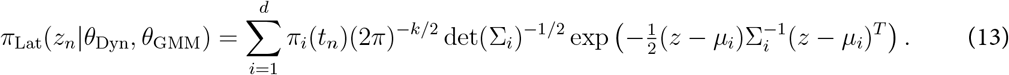

Here *t*_*n*_ is the time point that cell *n* was sampled. We assume that all latent vectors of cells *z*_*n*_ are independent. Hence, we ignore any potential dependencies due to common ancestry [59].

The likelihood function *L* consists of two factors

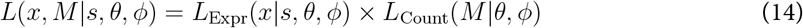

which are the likelihood of marker expression data *x*_*n*_, and the likelihood of cell counts *M*_*n*_, respectively. For the likelihood of the expression data, we start with a latent vector *z*_*n*_, and pass this through the decoder network *D*_*ϕ*_ to get parameters for the likelihood of *x*_*n*_:

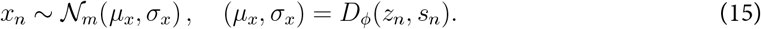

The likelihood of the count data is given by

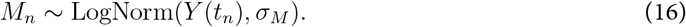

Finally, in order to perform inference, we have to specify a variational distribution *q*_*ϕ*_(*θ*). We assume that the variational distribution can be factorized as

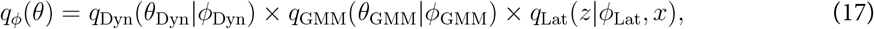

with factors for the dynamical model parameters, the GMM parameters and the latent vectors, respectively. For the (possibly transformed) dynamical model parameter vector

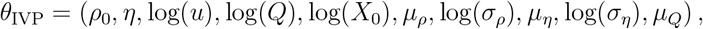

and *σ*_*M*_, we assume a multivariate normal and log-normal distribution, respectively:

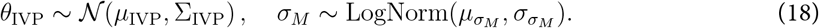

Next, we use a multivariate normal distribution for the location vectors of the mixture components;

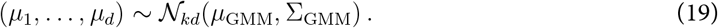

Finally, we have to specify a variational distribution for *z*. As we have a latent variable *z*_*n*_ for every observed marker expression vector *x*_*n*_, we use the method of amortization; instead of specifying variational parameter for each individual variable *z*_*n*_, we define a function *E* that calculates variational parameters *σ*_*z*_ and *µ*_*z*_, based on the data point *x*_*n*_:

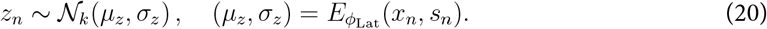

The function *E* is defined in terms of a neural network.

We implemented our method in the Pyro language for deep probabilistic programming [35]. We fitted the model to our data using stochastic variational inference (SVI). We used the Adam optimizer [60] with learning rate 2 *×* 10^−3^ and 10^5^ epochs, and used 20 parallel particles to evaluate the ∇ELBO. The neural network weights and biases were *L*_2_-regularized. All our code is available on https://github.com/chvandorp/scdynsys.git.

## Acknowledgments

This work was funded by the National Institutes of Health (NIAID), U01 AI150680 and R01 AI093870. We gratefully acknowledge Elise Bullock for technical support.

## Supporting Information

**Figure S1:**
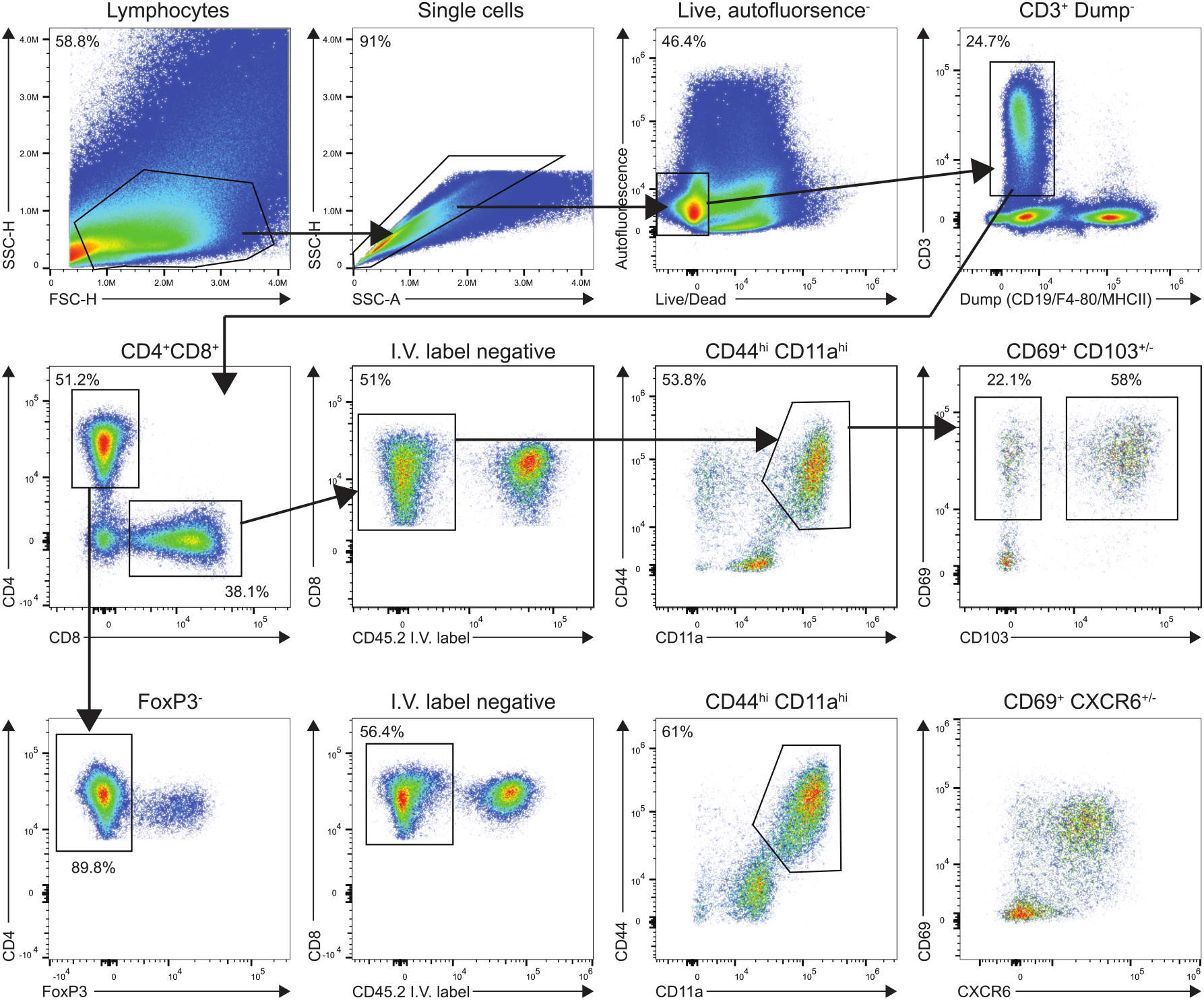
Gating Strategy for CD8+ and conventional CD4+ antigen experienced T cells residing in the lung.

**Figure S2:**
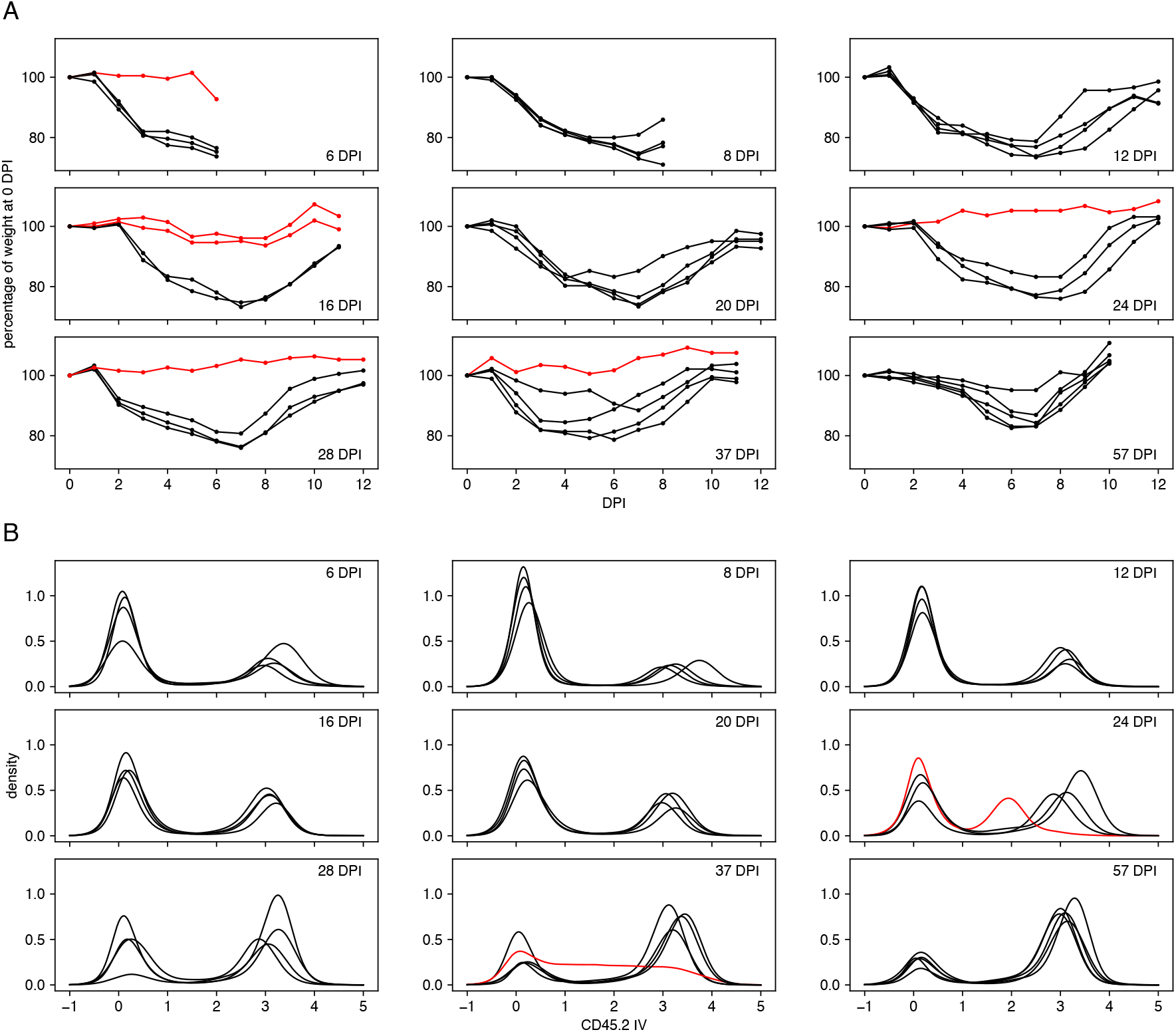
Weight curves of mice after IAV challenge and assessment of I.V. labeling. Each cohort contained 4 or 5 mice. In total data from 38 mice is presented. **A**. The curves indicate the percentage of the weight relative to that on the day of infection. The red curves correspond to mice that were excluded from further analysis due to lack of weight loss. **B**. Distribution of the IV label (CD45.2 IV) for each mouse. The red curves correspond to mice that were excluded from further analysis due to poor IV labeling.

**Figure S3:**
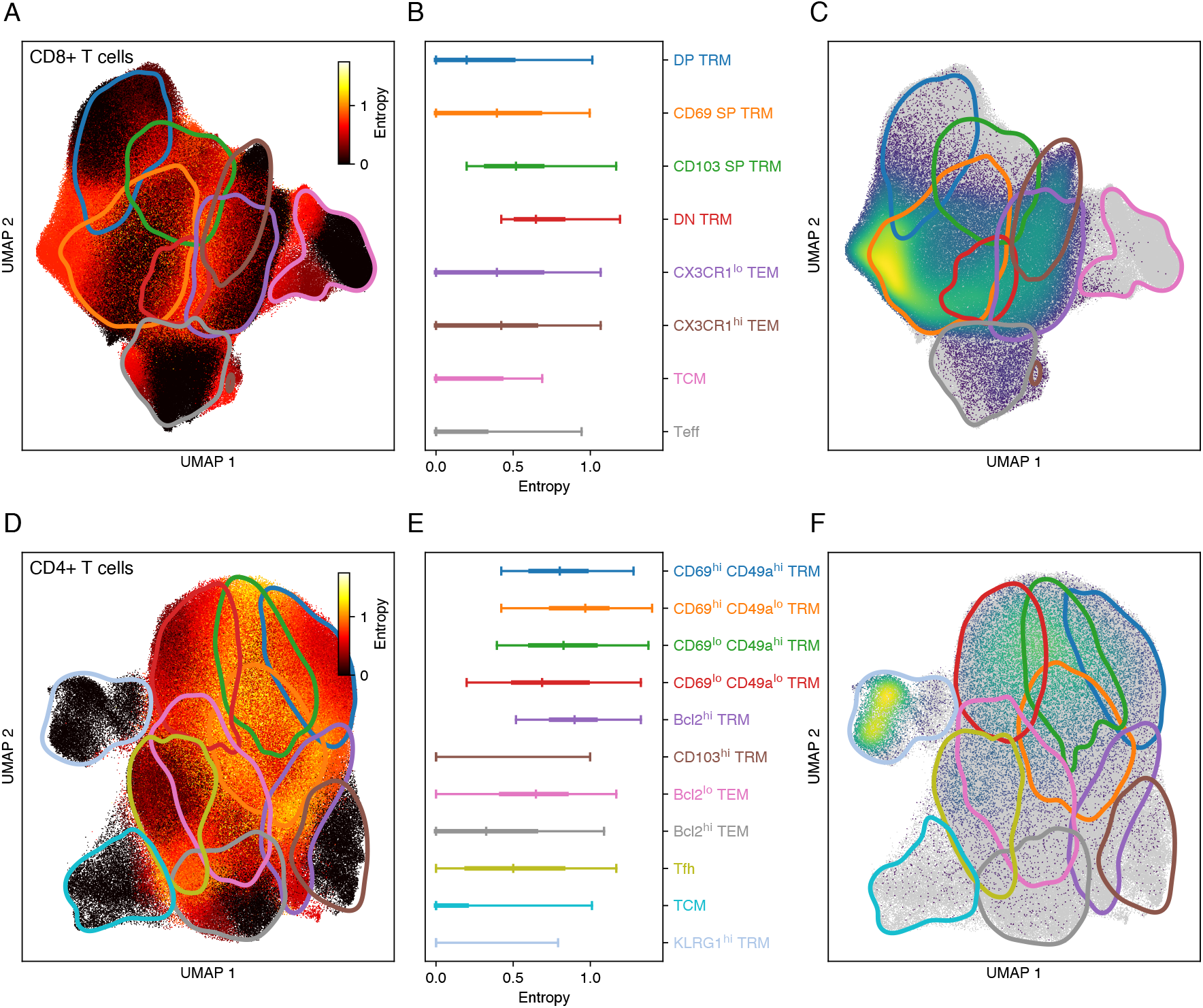
Validation of population assignment in the sequential approach. Results are based on data from *n* = 27 mice. **A**. Entropy of CD8 T cell population assignment based on 20 Leiden clustering runs with different random seeds. Black dots corresponds to very certain assignments, yellow dots to highly uncertain assignments. The colored contours indicate the location of the different sub-populations in the UMAP. **B**. Entropy distribution per cluster. The bar plots show the median, IQR and 2.5 - 97.5 percentile range. The color of the bars and labels correspond to the contours in panel A. **C**. Distribution of IAV NP-specific CD8 T cells in the UMAP with contours indicating clusters. Gray dots indicate bulk antigen-experienced cells. **D-F**. Same as panels A-C, but for CD4 T cell data.

**Figure S4:**
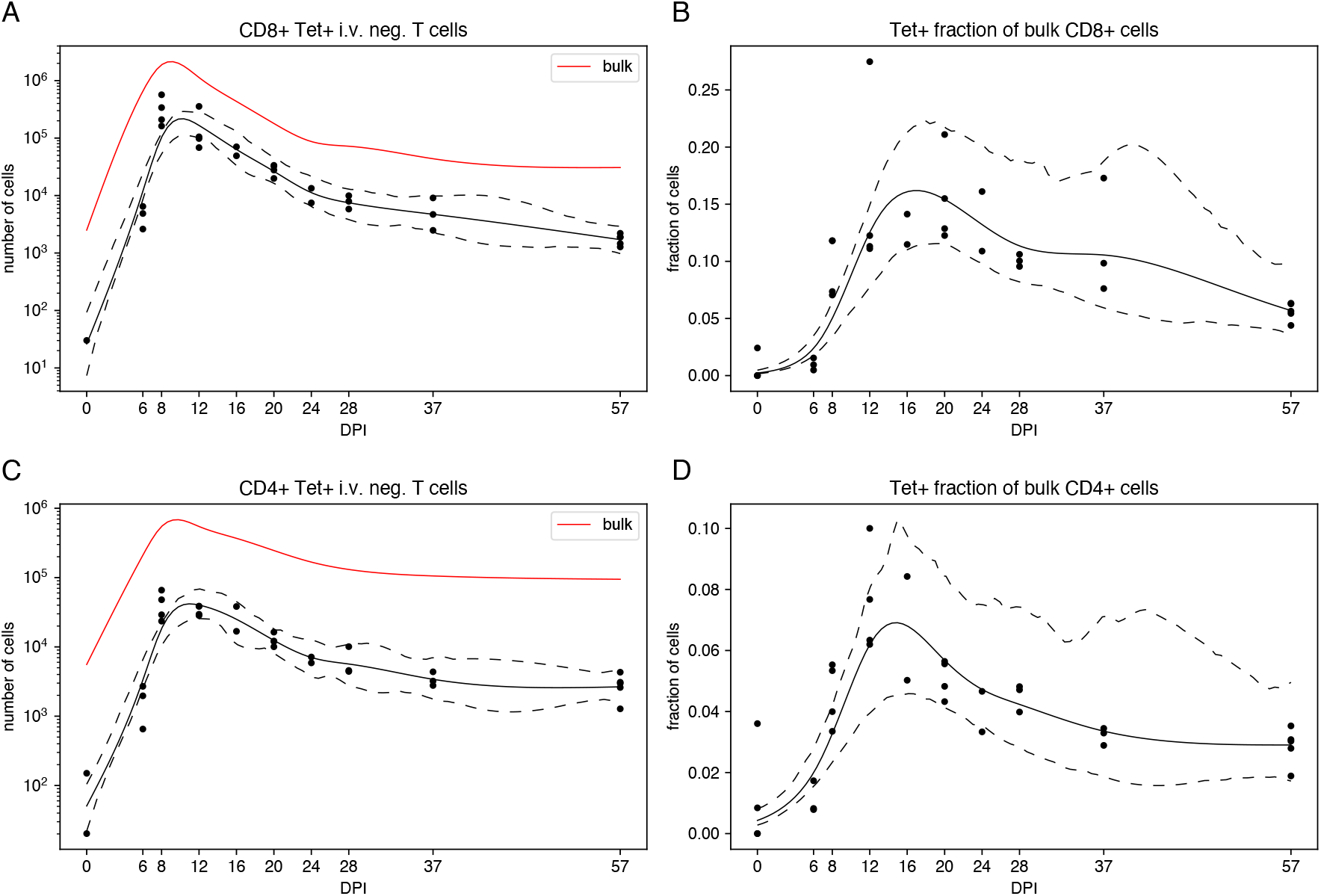
Timecourses of NP-specific T cells. Results are based on data from *n* = 34 mice. **A**. Number of NP-specific CD8 T cells, as a function of post-infection sampling time. The curve represents a spline fit to the log-transformed T cell counts, and the dashed lines represent the 95% confidence envelope (estimated by bootstrapping residuals). The red curve indicates the number of polyclonal CD8 T cells (cf. Fig. 1B). **B**. The Tet+ fraction of bulk CD8 T cells in the lung niche. Splines are fitted on the logit scale. **C**. Number of NP-specific CD4 T cells. **D**. The Tet+ fraction of bulk CD4 T cells.

**Figure S5:**
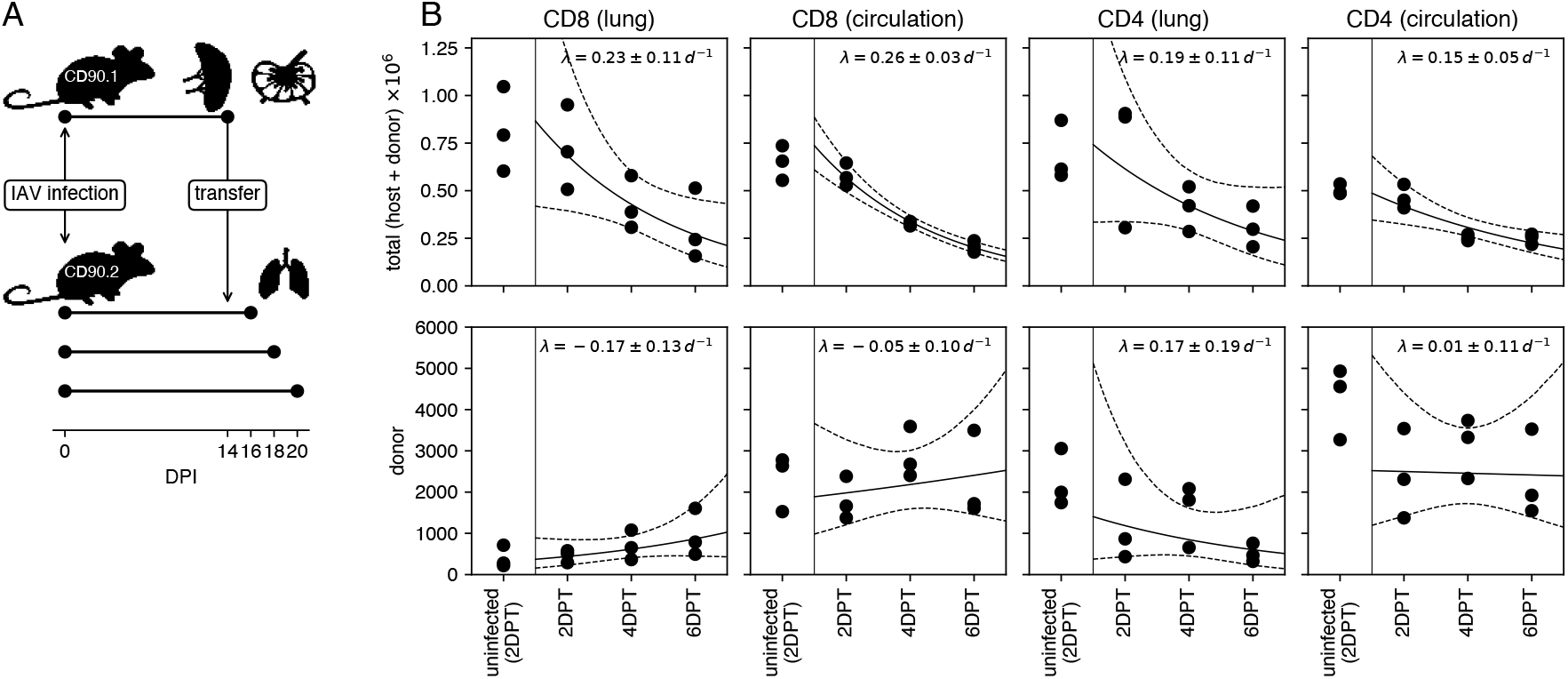
Congenic transfer experiment to assess the amount of ingress during the memory phase. Results are based on data from *n* = 12 mice in total, with 3 mice per group. **A**. Design of the congenic transfer experiment. **B**. Numbers of protected and labeled, antigen-experienced CD8 and CD4 T cells from host and donors combined (upper panels) and donors only (lower panels). We fitted a log-linear model to the cell counts (solid line: ML estimate, dashed lines: 95% confidence envelope). The indicated *λ* the estimated net loss rate (*±* standard error).

**Figure S6:**
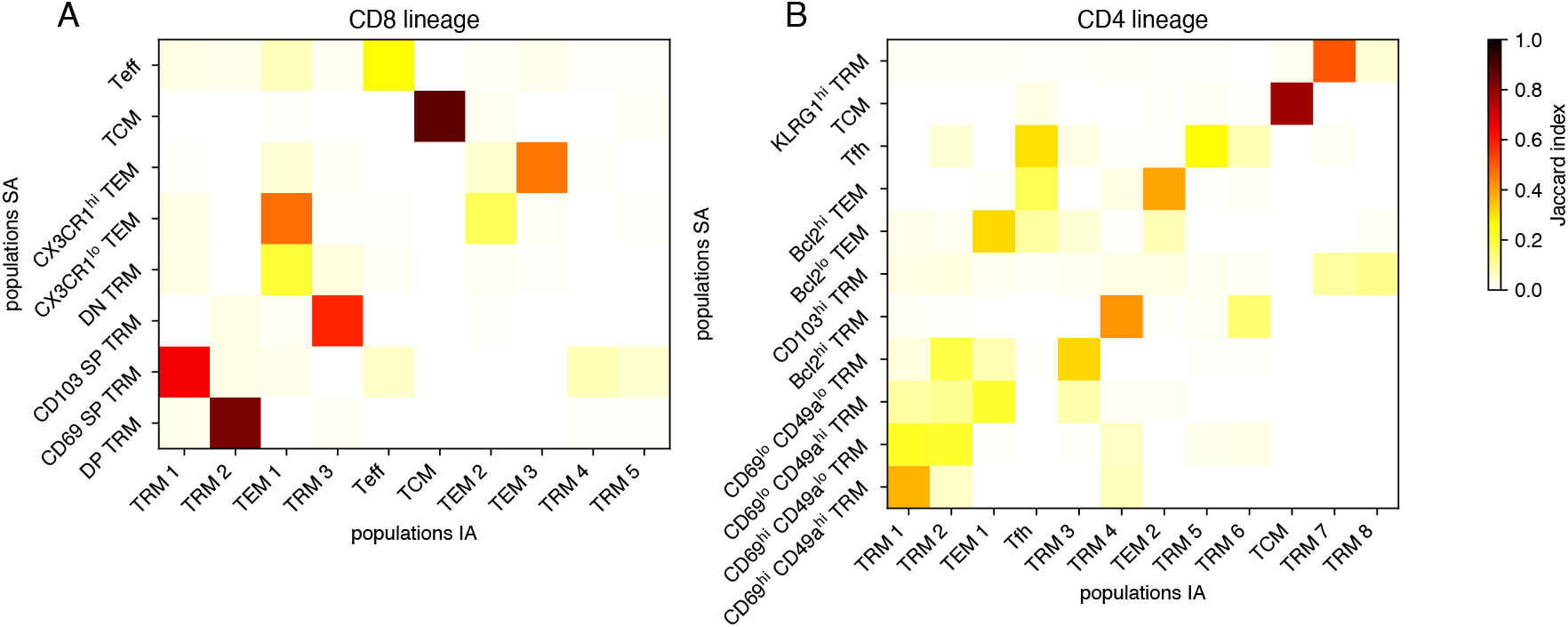
Comparison of results from the integrated and sequential approaches. Results are based on data from *n* = 27 mice. The sequential (SA) and integrated (IA) approaches both assign each cell to a subpopulation, but for any given cell this assignment may differ between the approaches. To quantify the similarity, we calculated the Jaccard index for each pair of sub-populations (one from the IA and the other from the SA). The Jaccard index is ratio of the number of cells that are in both clusters, and the number of cells that are in either one of the clusters. **A**. Results for CD8 T cell data. **B**. Results for CD4 T cell data.

**Figure S7:**
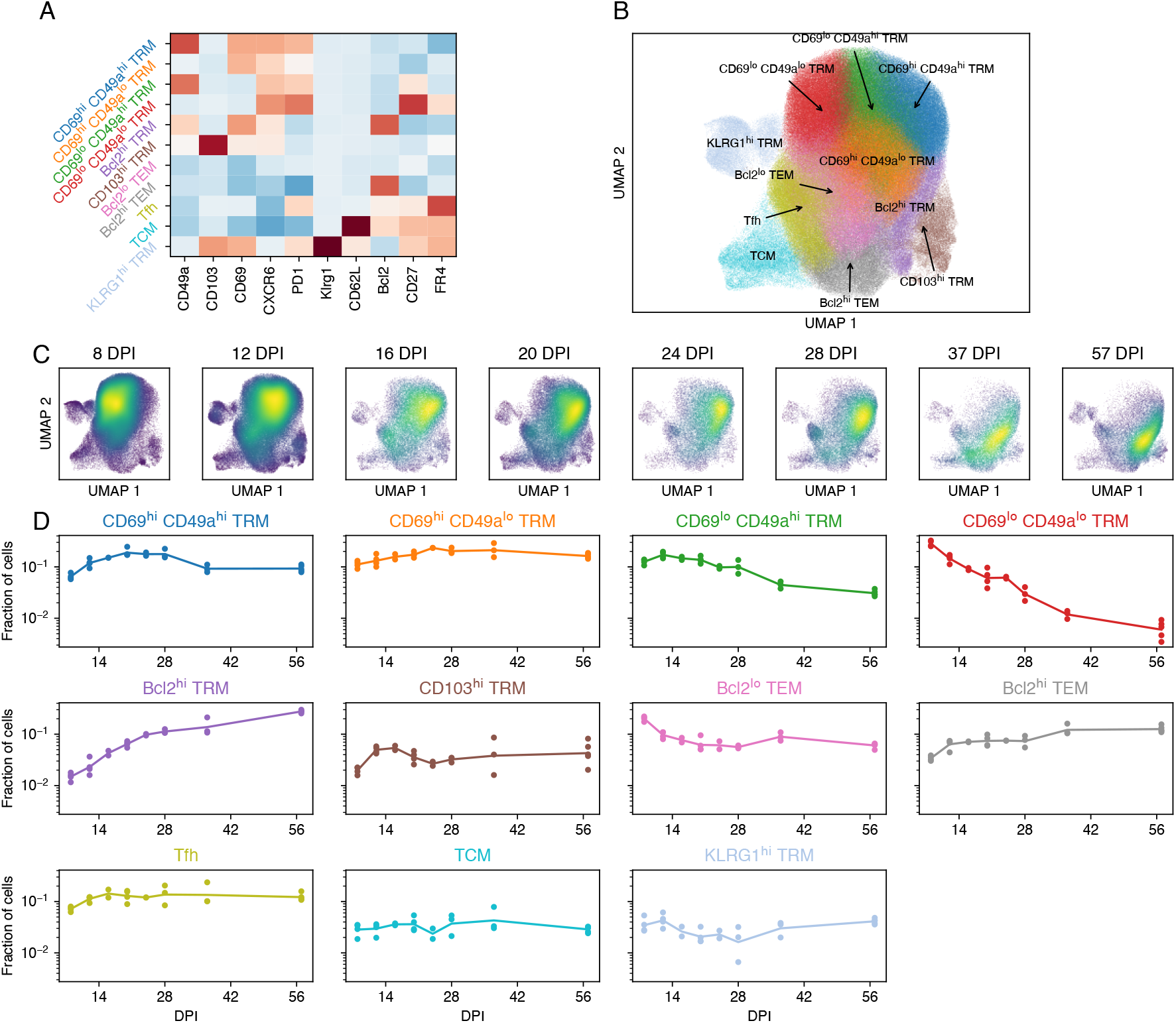
Pre-processing CD4+ T cell flow cytometry data for the sequential approach. Results are based on data from *n* = 27 mice. **A**. Marker expression heatmap for selected markers and consensus T-cell populations. **B**. UMAP of the marker expression data, colored by annotation. **C**. UMAPs of marker expression data, split by day post infection (DPI). The color scale reflects cell density in UMAP space. **D**. Time series of the fraction of cells in each cluster. The lines show a linear interpolation on the log scale.

**Figure S8:**
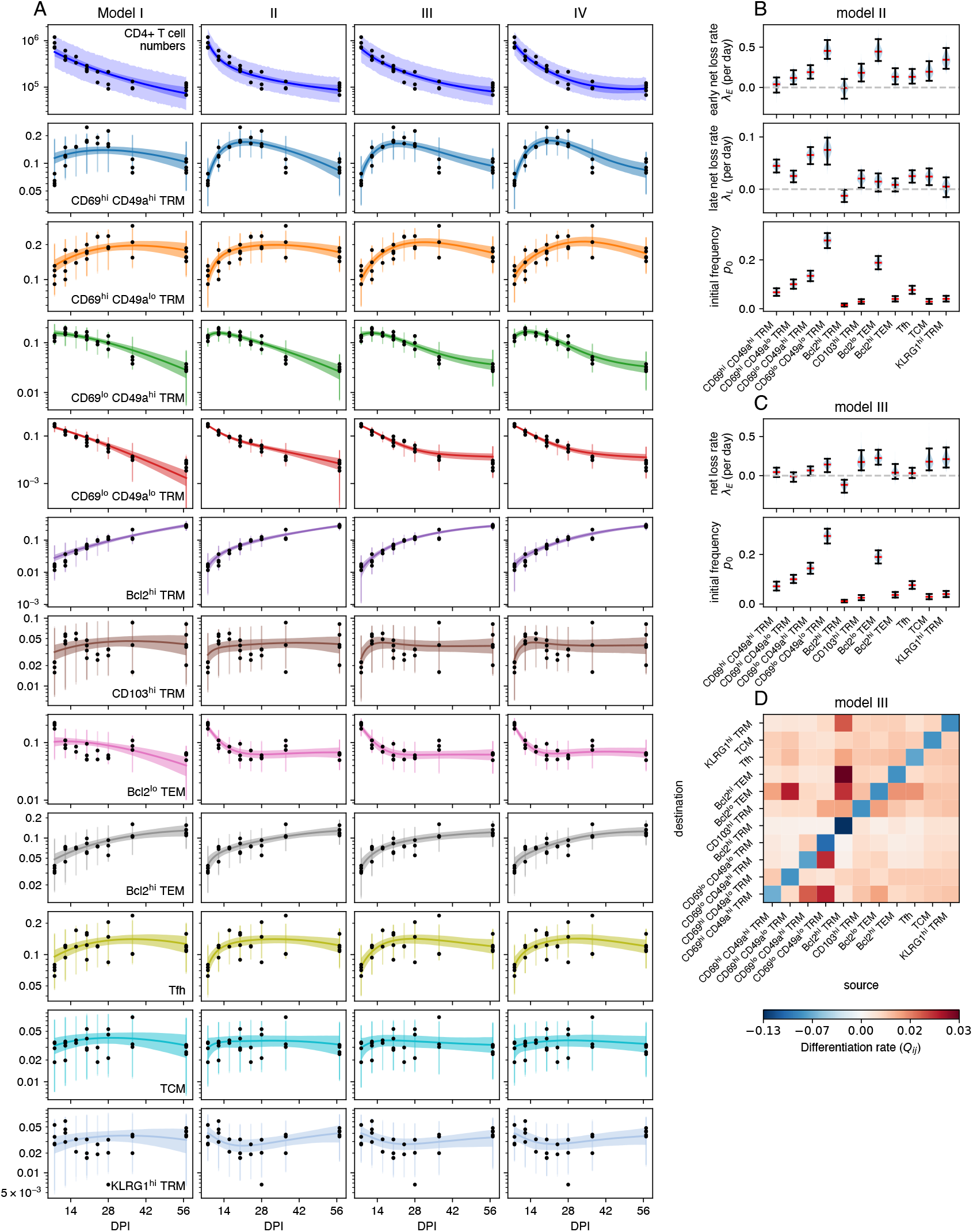
Model fits to CD4 T cell timeseries using the sequential approach. Results are based on data from *n* = 27 mice. **A**. Data and predictions from fitted models. Top panels show total antigen-experienced CD4 T cell counts in the lung, other panels show relative population sizes of each of the sub-populations. **B**. Parameter estimates using model II. **C**. Parameter estimates using model III. **D**. Estimated differentiation matrix in model III.

**Figure S9:**
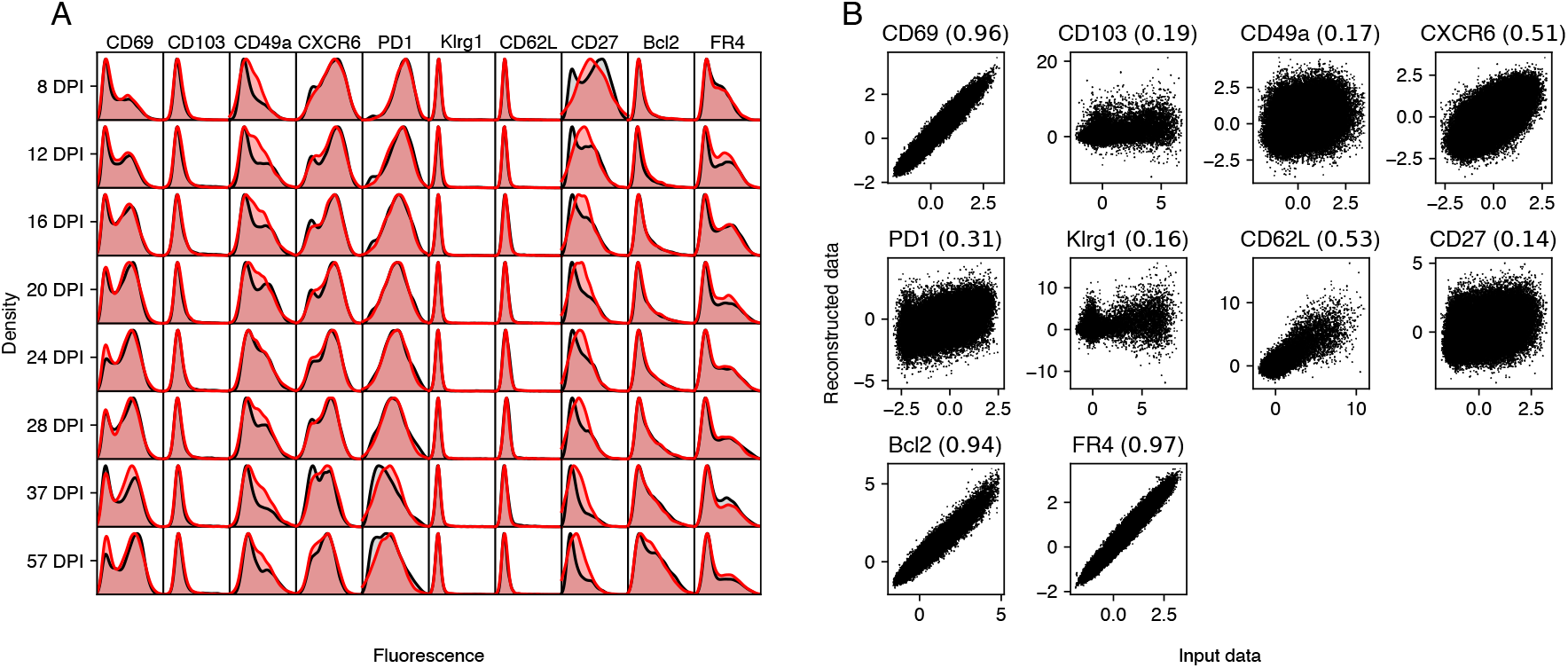
Posterior predictive checks for the integrated approach with CD4 T cell data. Results are based on data from *n* = 27 mice. **A**. Marginal distributions of marker expression (cf. Fig. 1, panel I). Data is shown in black, simulated data is shown in red. **B**. Input data and reconstruction using the auto-encoder model. The number in brackets is the coefficient of determination (*R*^2^).

**Figure S10:**
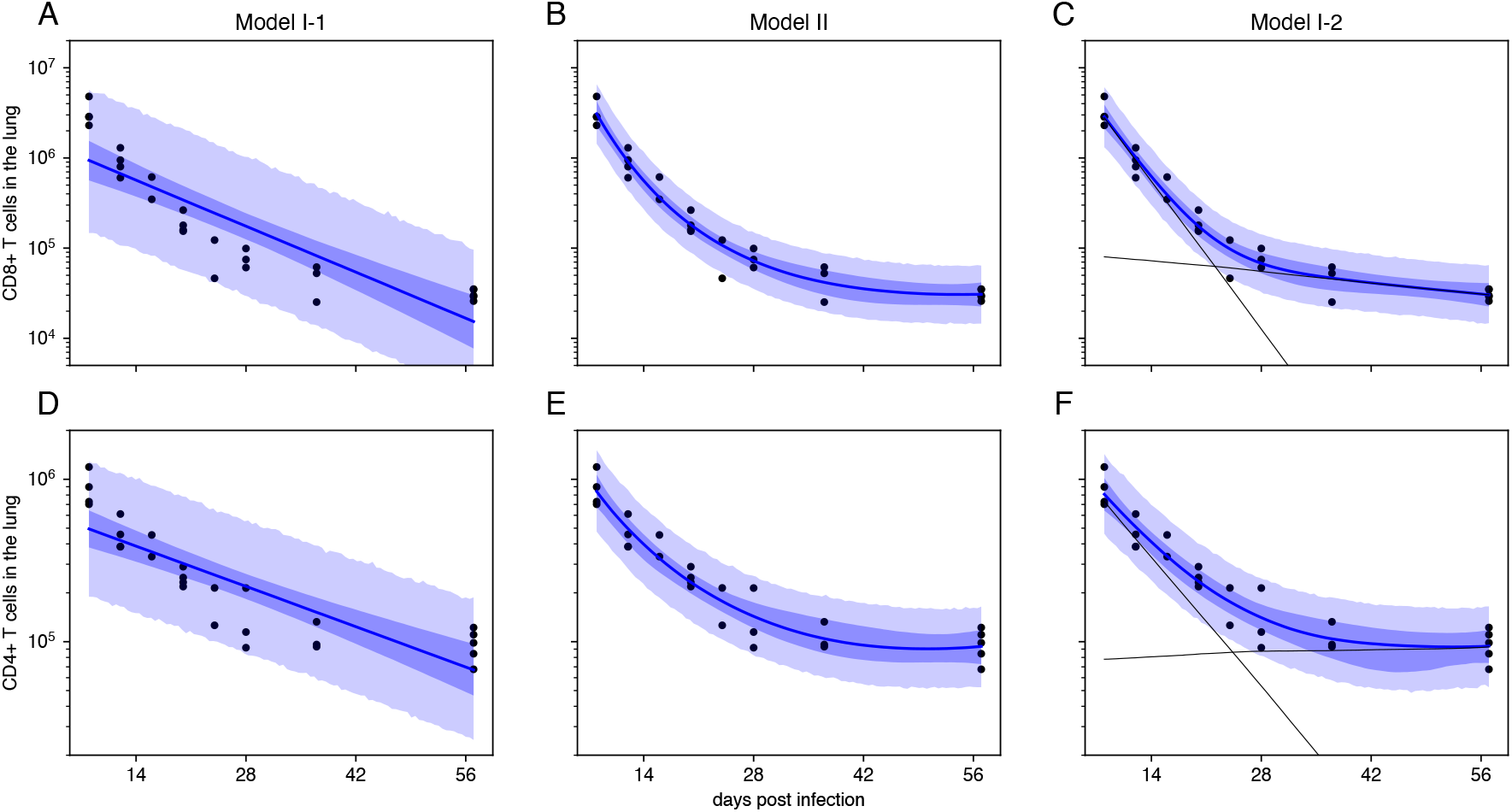
Model fits to cell count data alone. Results are based on data from *n* = 27 mice. **A**. time-homogeneous model with a single compartment (i.e. a log-linear model) fit to CD8 T-cell count data (dots). The model fit is shown as a blue curve (posterior median), with 95% CrI as a dark-blue band. The light-blue band shows the posterior predictive interval (i.e. simulated observations). **B**. Fit of model with a single compartment, but with time-dependent net loss rates *λ*(*t*). **C**. Fit of time-homogeneous model with two compartments. The population sizes (posterior median) of the two populations are shown as black curves. **D-F**. Fits of the three model to CD4 T cell counts.

**Figure S11:**
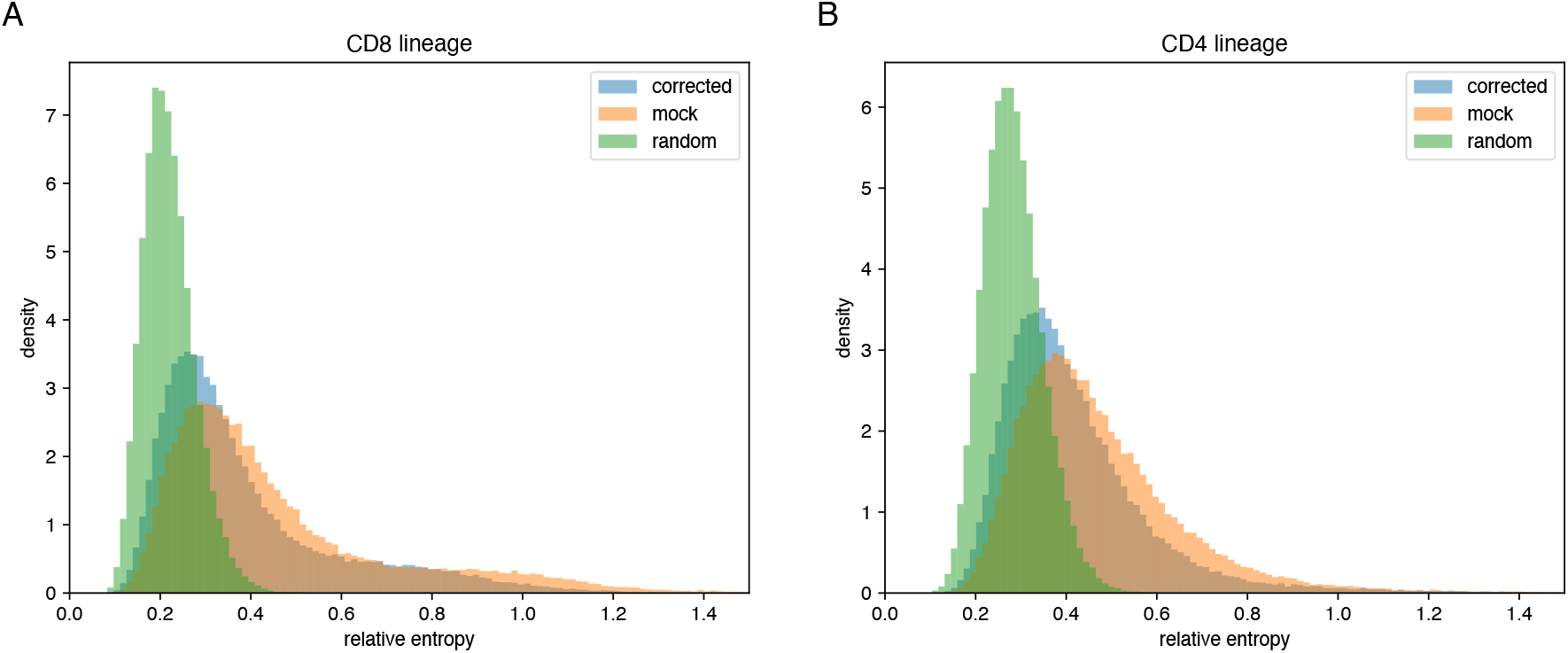
The effect of batch correction in the integrated approach. Results are based on data from *n* = 27 mice. Shown is the entropy of the experimental batch distribution around each cell, using the latent vector *z* and its nearest neighbors. Values for batch-corrected latent vectors are shown in blue. Mock corrected values are shown in orange, and values for randomized batch information are shown in green. Panels A and B show CD8 and CD4 data, respectively. The distributions are capped at 1.5 as there was a very small number of cells with high relative entropy.

**Figure S12:**
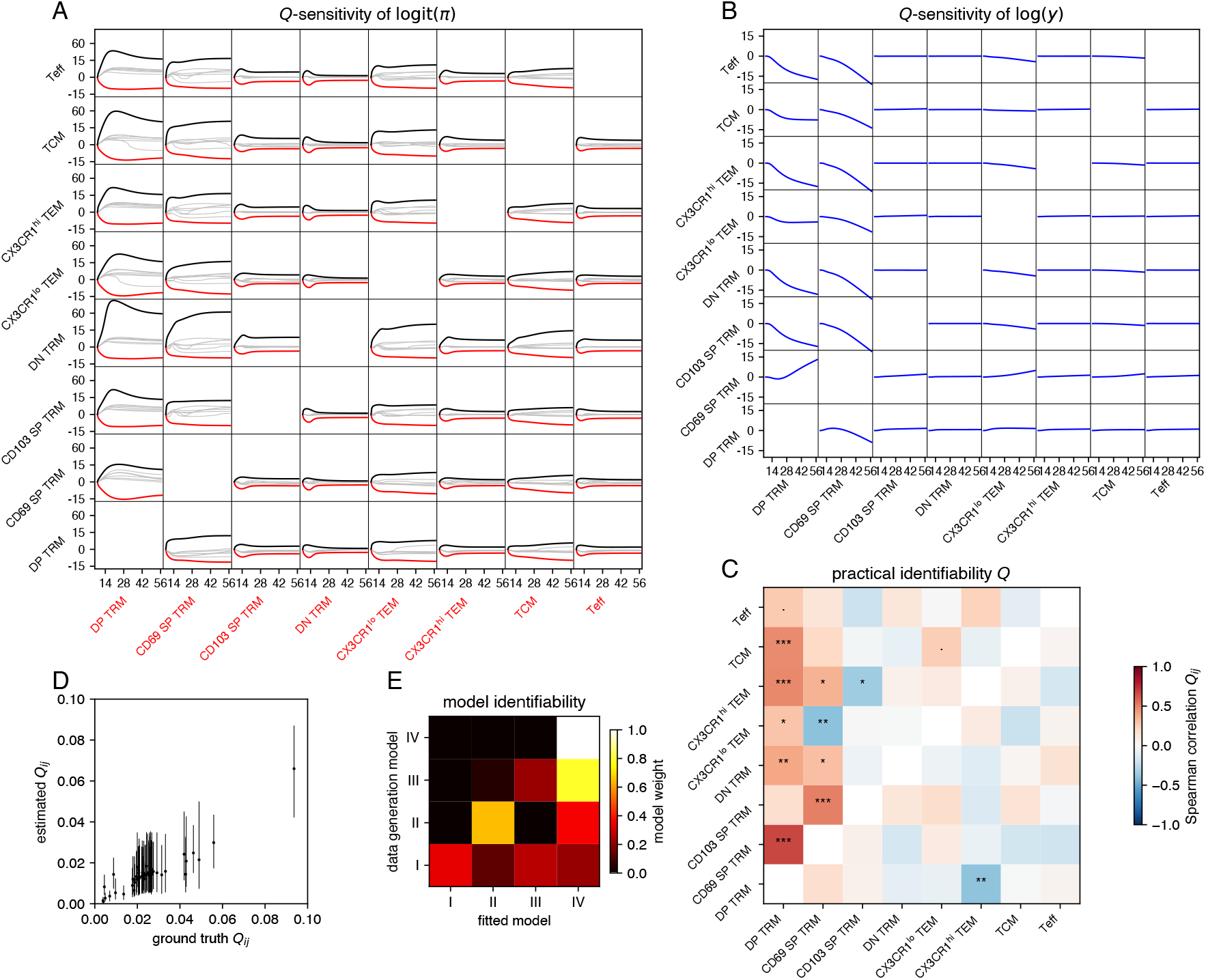
Identifiability and sensitivity analysis for the CD8 T cell lineage. **A**. Sensitivity of relative cluster sizes (*π*_*k*_(*t*)) with respect to differentiation rates *Q*_*ij*_. Each plot contains *d* = 8 curves *∂*logit(*π*_*k*_(*t*))*/∂Q*_*ij*_. The primary effects are highlighted in black (*i* = *k*) and red (*j* = *k*), while the secondary effects are shown in gray. The *y*-axes are shown on a squareroot-scale. **B**. Sensitivity of total population size (*Y* (*t*)) with respect to differentiation rates *Q*_*ij*_. The curves correspond to *∂* log(*Y* (*t*))*/∂Q*_*ij*_. **C**. Practical identifiability scores of differentiation rates *Q*_*ij*_. The heatmap shows the correlation coefficient between the ground truth value of *Q*_*ij*_, and the estimated value. The stars indicate the levels of statistical significance (· *p <* 0.1, ∗ *p <* 0.05, ∗∗ *p <* 0.01, ∗ ∗ ∗ *p <* 0.001) based on a range of 51 ground truth *Q*_*ij*_ values. **D**. A single pseudo-data set is simulated with model IV, using parameters estimated from the true data. Model IV is then fit to the pseudo-data, and for each pair of populations (*i, j*) we show the estimate versus the ground truth value of *Q*_*ij*_. **E**. Model identifiability. Data was simulated with and fit to each of the four models, resulting in 16 model fits. For a each simulated dataset, the four model fits are compared with model weights (shown in color). High diagonal model weights indicate that the ground-truth model is correctly identified. Shown is the median of 3 simulations for each model.

**Figure S13:**
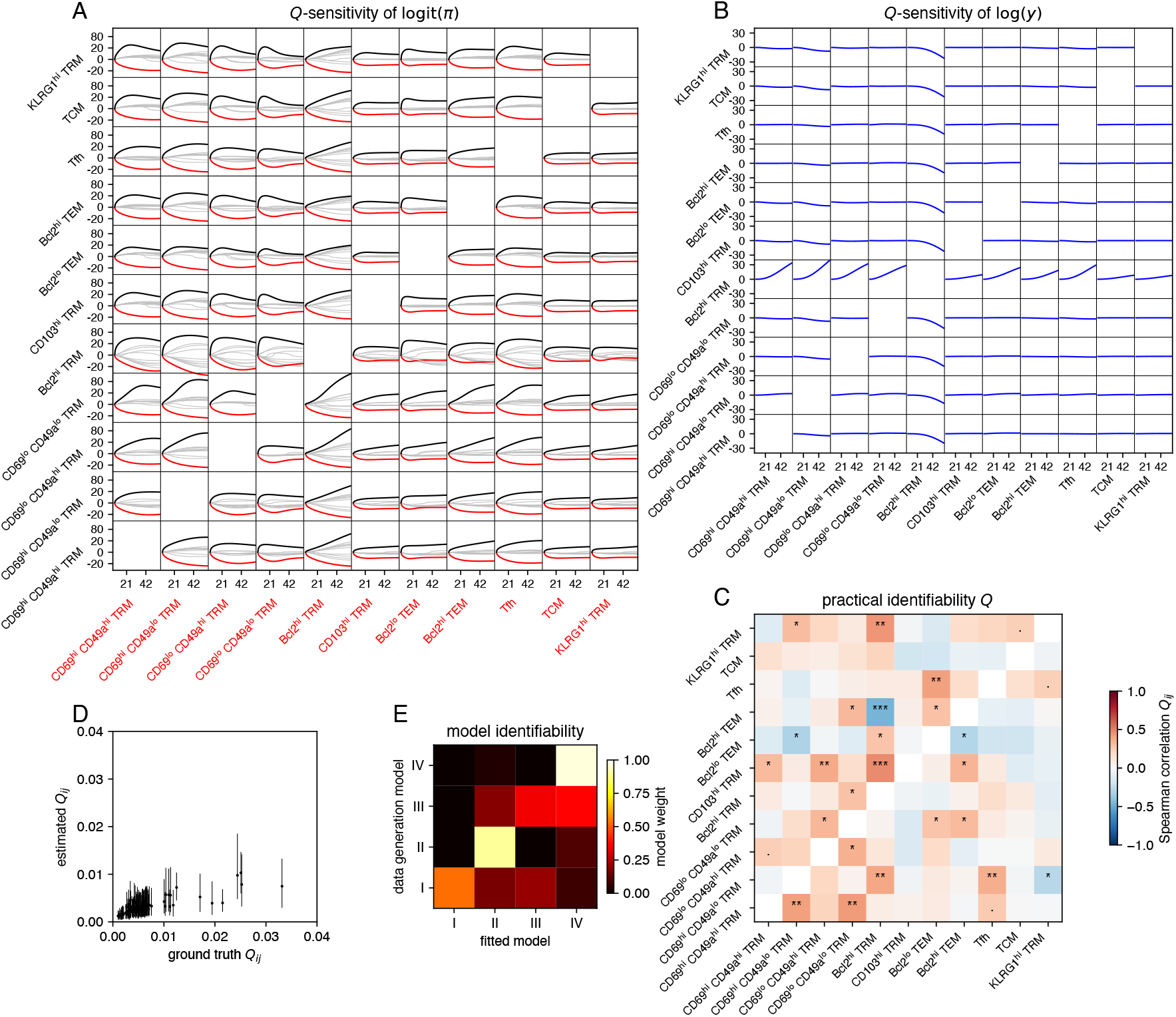
Identifiability and sensitivity analysis for the CD4 T cell lineage. See the caption of Fig. S12 for details. In this case we have *d* = 11 populations, and for panel D we simulated and fitted with model III.

**Figure S14:**
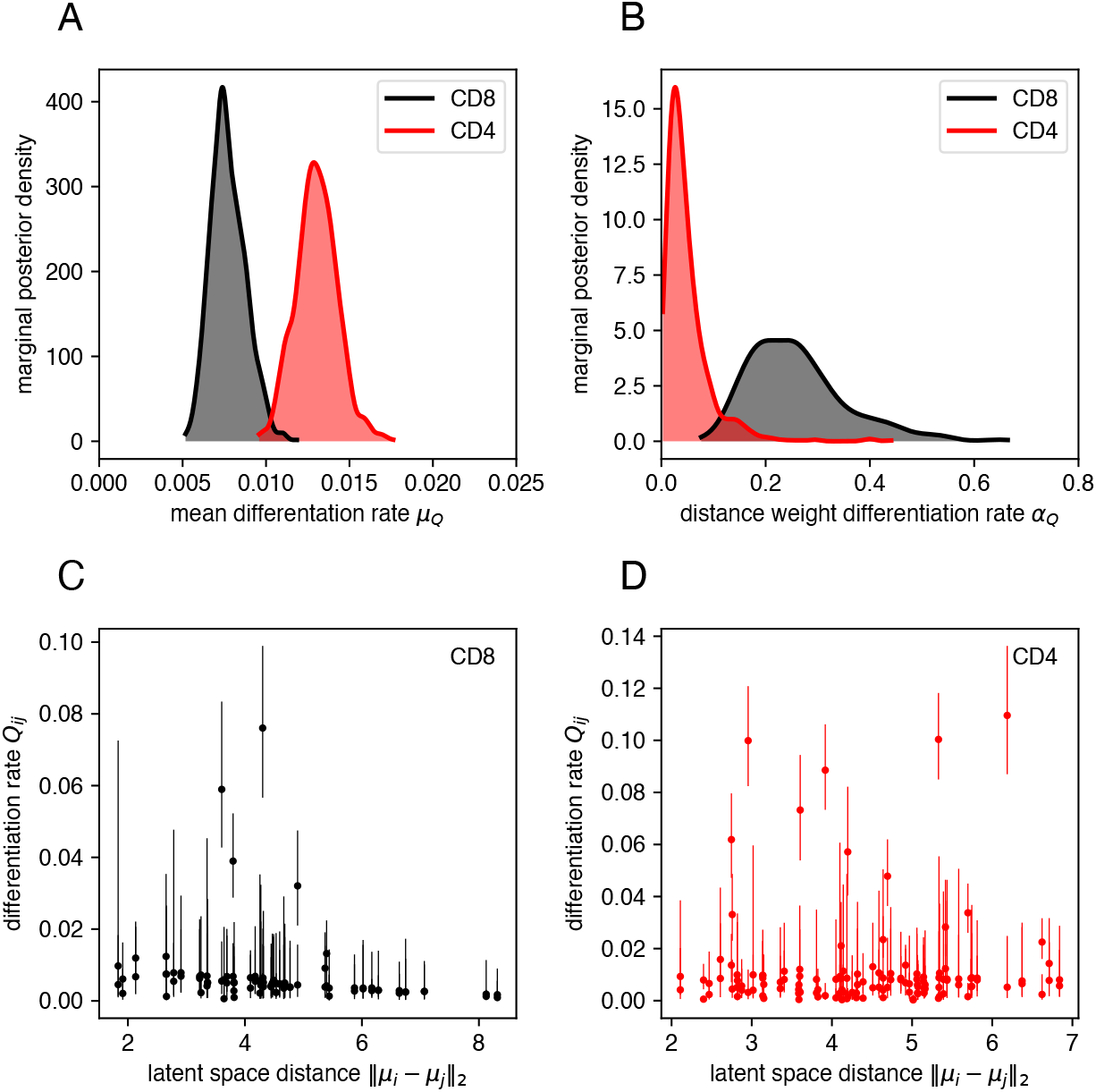
Cluster similarity informed prior distribution on the *Q*-matrix. **A**. Marginal posterior density of the mean differentiation rate *µ*_*Q*_. **B**. Marginal posterior density of the weight *α*_*Q*_ of the distance matrix *D*_*ij*_ = *‖ µ*_*i*_ − *µ*_*j*_ *‖* _2_ on the differentiation matrix elements *Q*_*ij*_. **C**. and **D**. 95% credible intervals (lines) and posterior medians (dots) of *Q*_*ij*_ as a function of the distance between the mixture components *i* and *j* in the latent space.

## SI Text A Ingress contributes minimally to the population dynamics of lung T_RM_ during the memory phase

To inform the construction of models of cell dynamics within the lung, we aimed to quantify the extent to which they were supplemented by new immigrants following the peak of infection. Our approach is illustrated in Fig. S5A. To do this we simultaneously infected cohorts of CD90.1 and CD90.2 congenic mice with IAV, and at day 14 post infection, transferred 106 cells from the spleens and mediastinal lymph nodes of the CD90.1 (donor) mice to the CD90.2 (host) mice (see Methods). At 16, 18 and 20 days post infection (2, 4, and 6 days post transfer, respectively), lung tissue was collected from the host mice, and analyzed with flow cytometry. The kinetics of accumulation of any donor cells in the host lung tissue would then reflect the degree of ongoing recruitment of new T_RM_ from circulating influenza-specific memory T cells.

Between 16 and 20 days post infection, the total numbers of CD4 and CD8 T cells (host+donor) fell by roughly 50% (Fig. S5B), consistent with the rates of decline observed in Fig. 1B-C. During the same time window, donor-derived cells were detectable in the lung but at low numbers that represented between 0.05-0.01% of transferred cells, and remained stable in number between 2-6 days post transfer. Between 16-18 DPI the rate of increase was 0.2 per day (95% CI [−0.1, 0.4]) for CD8+ T cells and −0.2 per day for CD4+ T cells (95% CI [−0.5, 0.2]). The number of donor cells in circulation (i.v. labeled cells from the lung sample) were consistently higher than cells in the tissue (CD8: 4.7 fold, *p* = 2 *×* 10^−4^; CD4: 3 fold, *p* = 4 *×* 10^−3^ paired t-test), while the total number of cells in circulation and in tissue were comparable (CD8: *p* = 0.13, CD4: *p* = 0.1). The number of donor cells in the lung tissue from uninfected donors was comparable to the number of cells from infected donors, indicating that the rate of ingress does not depend on IAV-specificity. We conclude that ingress of antigen-experienced T cells into the lung occurs at very low levels from at least 14 DPI onward.

## SI Text B A two-compartment, time-homogeneous model can explain the timecourse of CD8 T cell numbers

At first glance, the CD8 and CD4 T cell count data resemble typical biphasic exponential decay patterns [61]. Such time series can easily be modeled using two exponentially declining populations that decrease at two different rates. Initially, the more rapidly declining population is more abundant, but soon it is replaced (relatively) by the population that declines at a slower rate.

Because of this resemblance, it might be surprising that we need time-dependent loss rates or differentiation to model the T-cell populations in the lung. To make this initial expectation more rigorous, we fit three different models (denoted I-1, II and I-2) to the CD8 and CD4 T-cell count data alone. Model I-1 is time-homogeneous with only a single population (Fig S10A and D). This model can not describe the biphasic decay pattern seen in the data, and requires a large standard deviation for the error model (CD8: *σ*_*M*_ = 0.86, 95% CrI [0.67, 1.17]; CD4: *σ*_*M*_ = 0.45, 95% CrI [0.35, 0.61]).

Model II again has a single population, but now we include a time-dependent net loss rate *λ*(*t*) as in Eqn. 2. This model fits the T-cell count data very well (Fig S10B and E, and requires a much smaller standard deviation for the error model (CD8: *σ*_*M*_ = 0.34, 95% CrI [0.26, 0.46]; CD4: *σ*_*M*_ = 0.25, 95% CrI [0.19, 0.36]). Finally, model I-2 has constant decay rates, but assumes that there are two distinct T-cell populations that decay at different rates. Judging from the posterior predictive check (Fig. S10C and E), this model fits the data as well as model II, and requires similar *σ*_*M*_ values as model II.

In terms of LOO-IC, model II and I-2 are indistinguishable (CD8: ΔLOO-IC = 0.3 *±* 1.5; CD4: ΔLOO-IC = 0.3 *±* 0.5), while model I-1 is significantly worse (CD8: ΔLOO-IC = 24 *±* 3; CD4: ΔLOO-IC = 15 *±* 3). This means that model II and I-2 describe the count data equally well, and hence using count data alone, it is not possible to distinguish between a model with two populations or a time-dependent net loss rate.

## SI Text C The geometry of time-homogeneous loss of independent populations

The simplest model we considered has a geometric property that allows it to be easily tested against data. Suppose that the populations of T cells are independent (i.e. no differentiation, or *Q* = 0) and that their net loss rates are constant (i.e. time homogeneous). The model is then

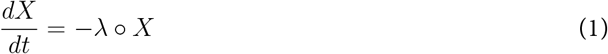

with initial condition *X*(*t*_0_) = *X*_0_. This model admits the following solution

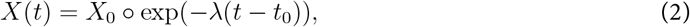

where the exponential function is taken element-wise. Again, we write 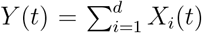 for the total population size and *π*_*i*_(*t*) = *X*_*i*_(*t*)*/Y* (*t*) for the population fractions.

When we look at the trajectories *π*_*i*_(*t*) on a logarithmic scale, a striking property is that they are all concave (Fig 3B and Fig. S8A, first column). We can see this mathematically as follows. A twice differentiable function is concave if the second derivative is non-positive. As the log of *π*_*i*_(*t*) is given by

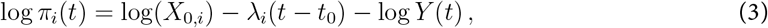

the second derivative is given by

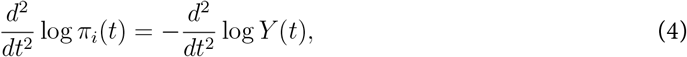

which does not depend on the population index *i*. The second derivative of log *Y* (*t*) is given by

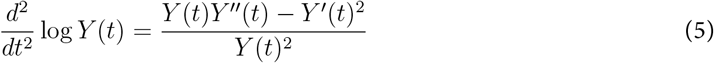

so its sign is determined by *Y* (*t*)*Y* ^*′′*^(*t*) − *Y* ^*′*^(*t*)^2^. We have 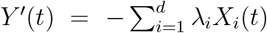, and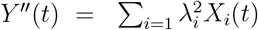. Now consider the vectors *a* and *b* given by 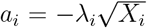 and 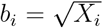. By the Cauchy-Schwartz inequality, we have *(a, b)*2 ≤ *(a, a)(b, b)*, and hence

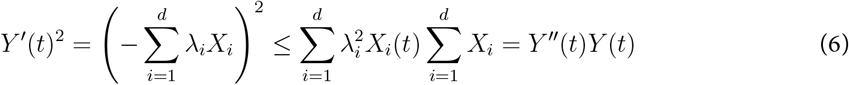

which means that 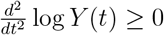. Hence log *π*_*i*_(*t*) is concave.

## SI Text D Batch correction

To measure how well batch correction performed in our integrated approach, we wanted to quantify the extent to which cells from different mice were well-mixed in phenotypic space [32]. To do this, we computed the K-nearest-neighbor graph of the latent representation *z*_*i*_ of the cells using the `scikit-learn` package [62]. For each cell *i*, we then counted the number of neighbors *k*_*i,s*_ derived from mouse *s* (including the focal cell). If cells are well mixed, all of these counts should be distributed according to the sample sizes *n*_*s*_ of the animals *s*. A convenient measure of how well the distributions of the *k*_*i,s*_ match is the relative entropy, given by

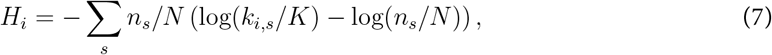

where *N* = ∑*s n*_*s*_ is the total number of cells from all animals. A lower relative entropy corresponds to more homogeneous mixing.

Fig. S11 shows the distribution of *H*_*i*_ as a blue histogram for the CD8 (panel A) and CD4 data (panel B). These distributions are difficult to interpret in isolation and so we compared them with two extremes. First, we computed the distribution of *H*_*i*_ under the assumption that all cells from all animals were homogeneously distributed. In this case we would have 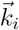 ~ Multinomial 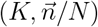. We randomly generated samples for each cell *i* and the resulting distribution is shown as an green histogram in Fig. S11. The other extreme is the case when we switch off the effect of batch-correction. To accomplish this, we picked the animal *s*^∗^ with the largest sample size *n*_*s*_∗ as a reference. We then used the encoder network to compute latent representations *z*_*i*_ from the pairs (*x*_*i*_, *s*^∗^), instead of the usual pairs (*x*_*i*_, *s*_*i*_). This means that we simulate the scenario in which all cells came from the same animal *s*^∗^. The resulting distribution of *H*_*i*_ is shown as an orange histogram in Fig. S11.

Our relative entropy-based analysis shows that batch correction is effective at aligning the distributions of cells from different animals within the latent space. However, the relative entropy remains much higher than expected under a perfectly homogeneous distribution (green vs. orange histogram in Fig. S11). This is due to the fact that mice are sampled at different DPI, and the phenotype distribution is dependent on time, and possibly because all data was collected at the same time (Fig. 1), reducing batch effects.

## SI Text E Sensitivity Analysis

To evaluate the sensitivity of ODE model (1) to the parameters, we derived and numerically integrated the sensitivity equations corresponding to the system. We therefore have to calculate the derivative of the state at each time point with respect to the parameter of interest. We can do this by interchanging the time derivative and the parameter gradient operator. Recall that the model is given by

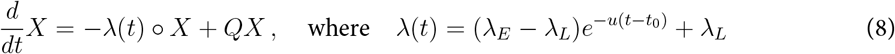

For completeness, we derive sensitivity equations for all parameters, altough we are mainly focussing on *Q* in the main text. The sensitivity equations for parameter *u* are then derived as follows.

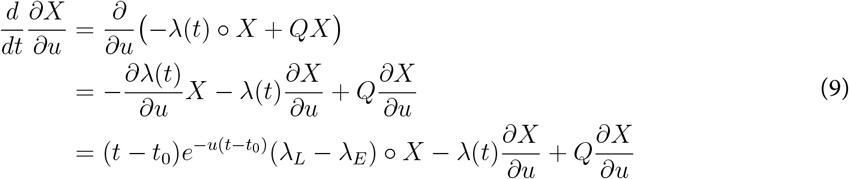

Next, we derive equations for the net loss rates. If we interpret ∂*/*∂*λ*_*E*_ as a row vector, ∂*X/*∂*λ*_*E*_ is a *d × d* matrix.

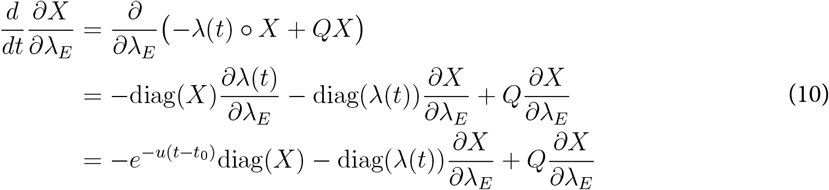

Likewise, we get for ∂*X/*∂*λ*_*L*_

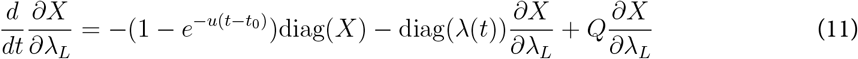

To derive sensitivity equations for the generator matrix *Q*, we first derive them for a general matrix *A* and the system 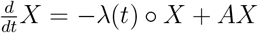, and then impose restrictions on the diagonal elements.

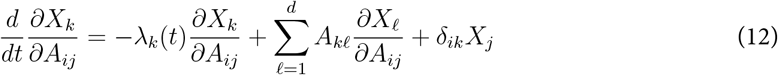

We now substitute *A*_*ij*_ = *Q*_*ij*_ such that 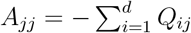 and get for *i≠ j*

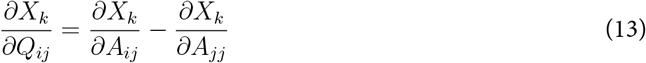

Therefore we get for *i≠ j*

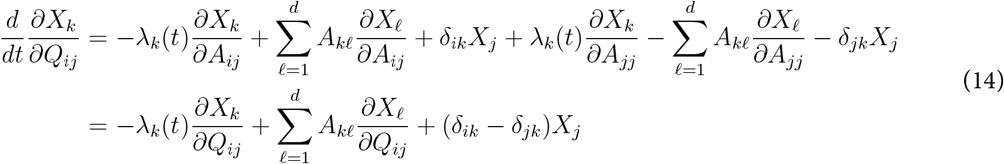

Finally, to compute the sensitivity of some transformation *f*(*X*) of the state *X* (e.g. *f*(*X*) = log(*Y*), or *f*(*X*) = logit(*π*_*i*_)), we simply apply the chain rule.

## Supporting Tables

**Table S1:** LOO-IC results for the sequential approach. Results are based on data from *n* = 27 mice. The models are ranked from best (top) to worst (bottom), using the “expected log predictive density” (elpd_loo) value. The p_loo value is a measure of the complexity of the model and generally increases with the number of parameters. The Δelpd value is the difference between the elpd_loo value and that of the best model. The se Δelpd is the standard error of the Δelpd.

| Model | elpd_loo | p_loo | $\Delta\text{elpd}$ | se $\Delta\text{elpd}$ |
| --- | --- | --- | --- | --- |
| *CD8 T-cell data* |  |  |  |  |
| IV | -1215.5 | 29.1 | - | - |
| II | -1253.9 | 24.3 | 38.4 | 9.6 |
| III | -1288.4 | 19.3 | 72.9 | 13.2 |
| I | -1372.9 | 22.2 | 157.4 | 29.9 |
| *CD4 T-cell data* |  |  |  |  |
| IV | -1805.4 | 39.0 | - | - |
| II | -1812.1 | 32.5 | 6.7 | 5.1 |
| III | -1816.4 | 33.9 | 11.0 | 4.9 |
| I | -1889.7 | 24.9 | 84.3 | 21.2 |

**Table S2:**
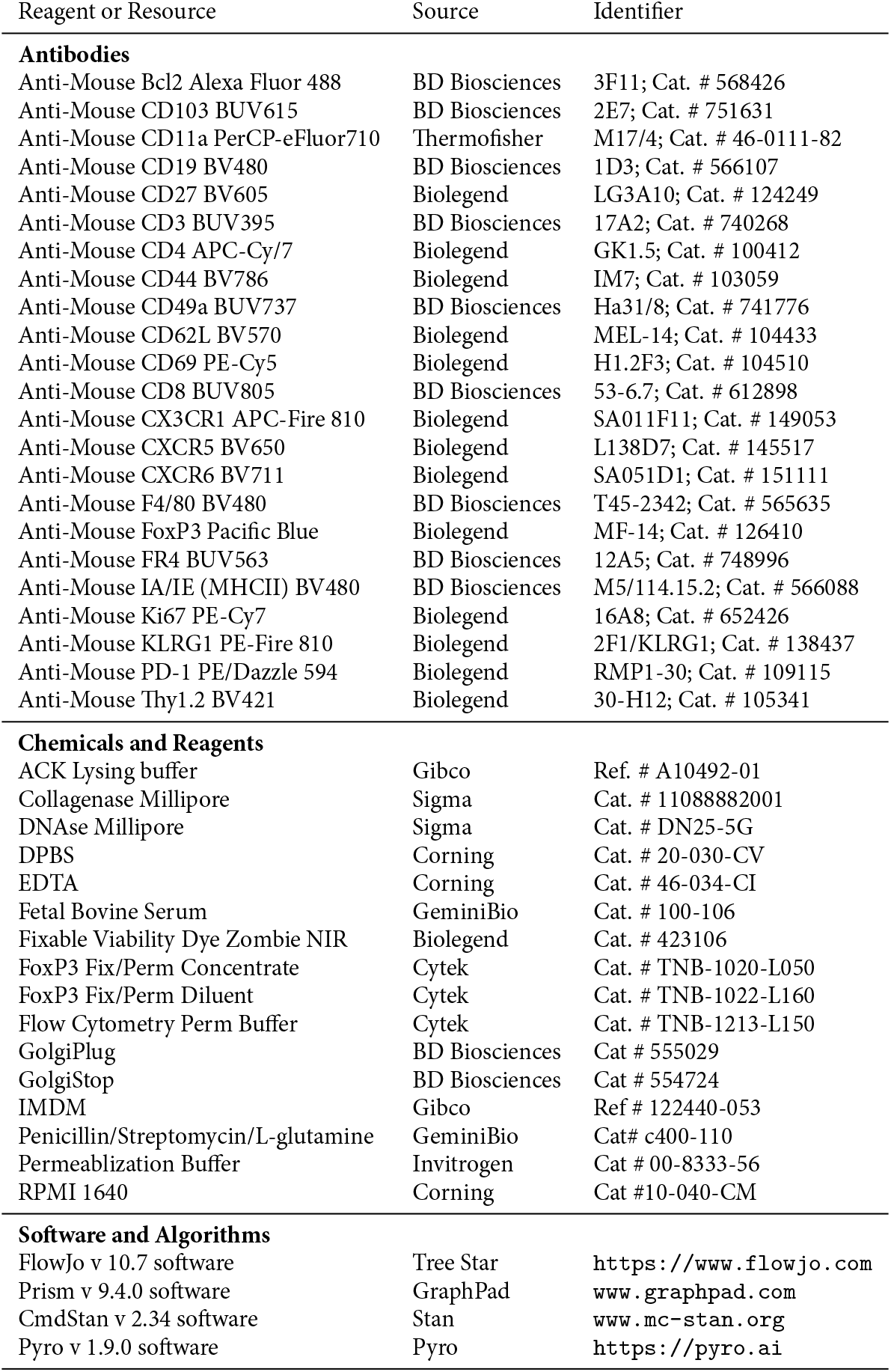
Reagents and resources.

**Table S3:** Prior distributions for the Bayesian model. Normal distributions are parameterized with scale parameters instead of variance. The half-normal distribution is denoted HalfNorm(*σ*), and the log-normal distribution LogNorm(*µ, σ*). The prior for *Q*_*ij*_ in the integrated approach is informed by the distance *D*_*ij*_ = *‖ µ*_*i*_ − *µ*_*j*_ *‖* _2_ between GMM cmponent *i* and *j* in the latent space.

| Parameter | Description | Prior | Hyper-parameters |
| --- | --- | --- | --- |
| $\lambda_{E,i}$ | Initial net loss rate of cluster $i$ | $\lambda_{E,i} \sim \mathcal{N}(\mu_{\lambda_E}, \sigma_{\lambda_E})$ | $\mu_{\lambda_E} \sim \mathcal{N}(0, 1), \sigma_{\lambda_E} \sim \text{HalfNorm}(1)$ |
| $\lambda_{L,i}$ | Long-term net loss rate of cluster $i$ | $\lambda_{L,i} \sim \mathcal{N}(\mu_{\lambda_L}, \sigma_{\lambda_L})$ | $\mu_{\lambda_L} \sim \mathcal{N}(0, 1), \sigma_{\lambda_L} \sim \text{HalfNorm}(1)$ |
| $u$ | Rate at which net loss rate $\lambda$ changes from $\lambda_E$ to $\lambda_L$ | $u \sim \text{HalfNorm}(1)$ | - |
| $Q_{ij}$ | Rate of differentiation from cluster $j$ to $i$ | $Q_{ij} \sim \text{Exp}(\mu_Q^{-1})$ | $\mu_Q \sim \text{Exp}(100)$ |
| $X_{0,i}$ | Initial size of cluster $i$ | $X_{0,i} \sim \text{LogNorm}(0, 10)$ | - |
| $\sigma_M$ | Scale parameter for likelihood of T cell numbers | $\sigma_M \sim \text{Exp}(1)$ | - |
| $\tau_K$ | Dispersion parameter for likelihood of cluster size | $\tau_K^{-1} \sim \text{Exp}(10^3)$ | - |
| *Specialized priors for the integrated approach* |  |  |  |
| $Q_{ij}$ | Rate of differentiation from cluster $j$ to $i$ | $Q_{ij} \sim \text{Exp}(\mu_Q^{-1} \exp(\alpha_Q(D_{ij} - \bar{D})))$ | $\mu_Q \sim \text{Exp}(100), \alpha_Q \sim \text{HalfNorm}(1)$ |

